# Pan-gastrointestinal prioritization links *LINC01094* to a colorectal cancer transcriptional state and replication-associated checkpoint activation

**DOI:** 10.64898/2026.05.26.728026

**Authors:** Bridget Flowers, Penelope Lialios, Eileen Park, Nathan Smith, Rebecca J. Burgess, Justin Whalley, Jong-Sun Lee

## Abstract

Cancer transcriptomes nominate many disease-associated long noncoding RNAs (lncRNAs), but functional prioritization remains difficult. Here, we integrated recurrent tumor upregulation, clinical association, co-expression network context, and malignant epithelial detectability across colorectal, gastric, and esophageal cancers, identifying five lncRNAs with convergent clinical and network support. We selected the intergenic lncRNA *LINC01094* for functional characterization in colorectal cancer (CRC). Across 52 CRC cell lines, endogenous *LINC01094* expression defined a transcriptional state linking higher expression to extracellular matrix, adhesion, and inflammatory programs and lower expression to DNA replication, mitochondrial, and ribosome/RNP programs. Acute CRISPR interference (CRISPRi)-mediated depletion in HCT116 cells produced a nearly inverse transcriptional response; among 764 GO terms significant in both analyses, 748 (97.9%) changed in opposite directions (Pearson *r* = −0.962). Functionally, *LINC01094* depletion impaired cell growth in both HCT116 and Caco-2 cells. In HCT116 cells, depletion reduced DNA synthesis despite induction of replication-associated transcriptional programs and was accompanied by increased γH2AX, CHK1 Ser345 phosphorylation, and p21 accumulation without detectable apoptotic execution. γH2AX foci were also elevated within residual EdU-positive nuclei. The predominant *LINC01094* transcript in HCT116 cells was nuclear-enriched, whereas *SLPI*, a reported effector of a cytoplasmic miR-1266-5p/SLPI ceRNA model in CRC, was not reproducibly altered following *LINC01094* depletion in either CRC model. These findings link *LINC01094* to maintenance of a CRC-associated transcriptional state and show that its acute disruption is accompanied by impaired productive DNA synthesis, replication-associated DNA-damage signaling, and checkpoint activation, supporting a broader, context-dependent role for LINC01094 beyond previously described pathway-specific models.

## INTRODUCTION

Gastrointestinal (GI) cancers arise from distinct epithelial lineages across the esophagus, stomach, colon, and rectum, yet share recurrent tumor-associated molecular features and collectively account for substantial cancer mortality (1). Pan-cancer and GI-focused studies have shown that these tumors nonetheless retain strong tissue-of-origin and lineage-specific transcriptional identities (1–3), and single-cell and spatial transcriptomic studies have revealed further heterogeneity among the malignant epithelial, stromal, immune, and endothelial populations within individual tumors (4, 5). These two forms of heterogeneity complicate identification of recurrent tumor-associated signals across GI cancers.

Long noncoding RNAs (lncRNAs) comprise a large and incompletely annotated class of regulatory transcripts with roles in transcription, RNA processing, nuclear organization, signaling, stress responses, and translational control (6, 7). LncRNA dysregulation has been linked to tumor growth, metastasis, and adverse clinical outcomes in GI malignancies (8–11). Yet, systematic prioritization of lncRNAs for functional interrogation remains difficult. Unlike protein-coding genes, lncRNAs often lack defined functional domains and established pathway annotations (6, 7, 12, 13), and tissue-, lineage-, and context-dependent expression can confound disease-relevant signals with anatomical origin and tumor composition (7, 14–17). The central challenge is therefore to identify which disease-associated lncRNAs warrant direct functional investigation.

The Cancer Genome Atlas (TCGA) provides genome-wide expression profiles from matched patient tumor and normal tissue together with clinicopathologic annotations, enabling identification of recurrent tumor-associated lncRNAs and correlations with disease progression and/or patient survival (2, 3). Co-expression network analysis places lncRNAs within disease-associated transcriptional programs and measures their connectivity within those programs (18, 19). Bulk tumor profiles, however, cannot assign lncRNA expression to the malignant compartment (16, 17). Single-cell transcriptomic data enable candidate lncRNA detection within distinct cell populations (4). Each independent analysis can generate an extensive catalog of disease-associated lncRNAs for which selection and functional testing of relevant candidates remains difficult and experimentally demanding.

Here, we evaluated recurrent tumor upregulation, clinical association, co-expression network context, and malignant epithelial detectability independently across colorectal cancer (CRC), stomach adenocarcinoma (STAD), and esophageal carcinoma (ESCA), integrating these dimensions only during candidate selection. This identified five lncRNAs with clinical and network support and prioritized the intergenic lncRNA *LINC01094* for functional characterization in CRC. Previous work has linked *LINC01094* to CRC progression and proliferation and proposed a cytoplasmic microRNA-dependent regulatory model (20), but whether its endogenous expression marks a broader CRC transcriptional state and how that state responds to acute *LINC01094* loss remained unclear. Across 52 CRC cell lines, endogenous *LINC01094* expression linked high expression to ECM/adhesion and inflammatory programs and low expression to replication, mitochondrial, and ribosome/RNP programs. Acute CRISPRi-mediated depletion in HCT116 cells shifted the transcriptome toward the low-*LINC01094* state while reducing DNA synthesis and inducing replication-associated DNA-damage signaling and checkpoint activation. Together, these findings support a role for *LINC01094* in maintaining a CRC-associated transcriptional state and demonstrate how multidimensional disease evidence can guide functional investigation of lncRNAs.

## RESULTS

### Pan-GI cancers exhibit lncRNA dysregulation

Because lncRNA expression is particularly tissue- and context-dependent (14, 15), we reasoned that recurrent changes across GI cancers would more likely reflect shared tumor biology than lineage identity. We therefore evaluated recurrent tumor upregulation, clinical association, co-expression network context, and malignant epithelial detectability separately because they address distinct biological questions and limitations, integrating their outputs only at candidate selection (Fig. 1A). We first examined global expression patterns across CRC, STAD, and ESCA, three anatomically related GI epithelial malignancies with transcriptomic and clinical annotation available for matched tumor–normal and cross-cancer analysis (Table S1A). Principal component analysis (PCA) broadly separated tumor from matched normal tissues, while tumors partially overlapped and retained cancer-type-associated identities (Figs. 1B and S1A). These results established tumor status as a major source of transcriptional variation across the datasets while preserving tissue-of-origin differences.

**Figure 1.**
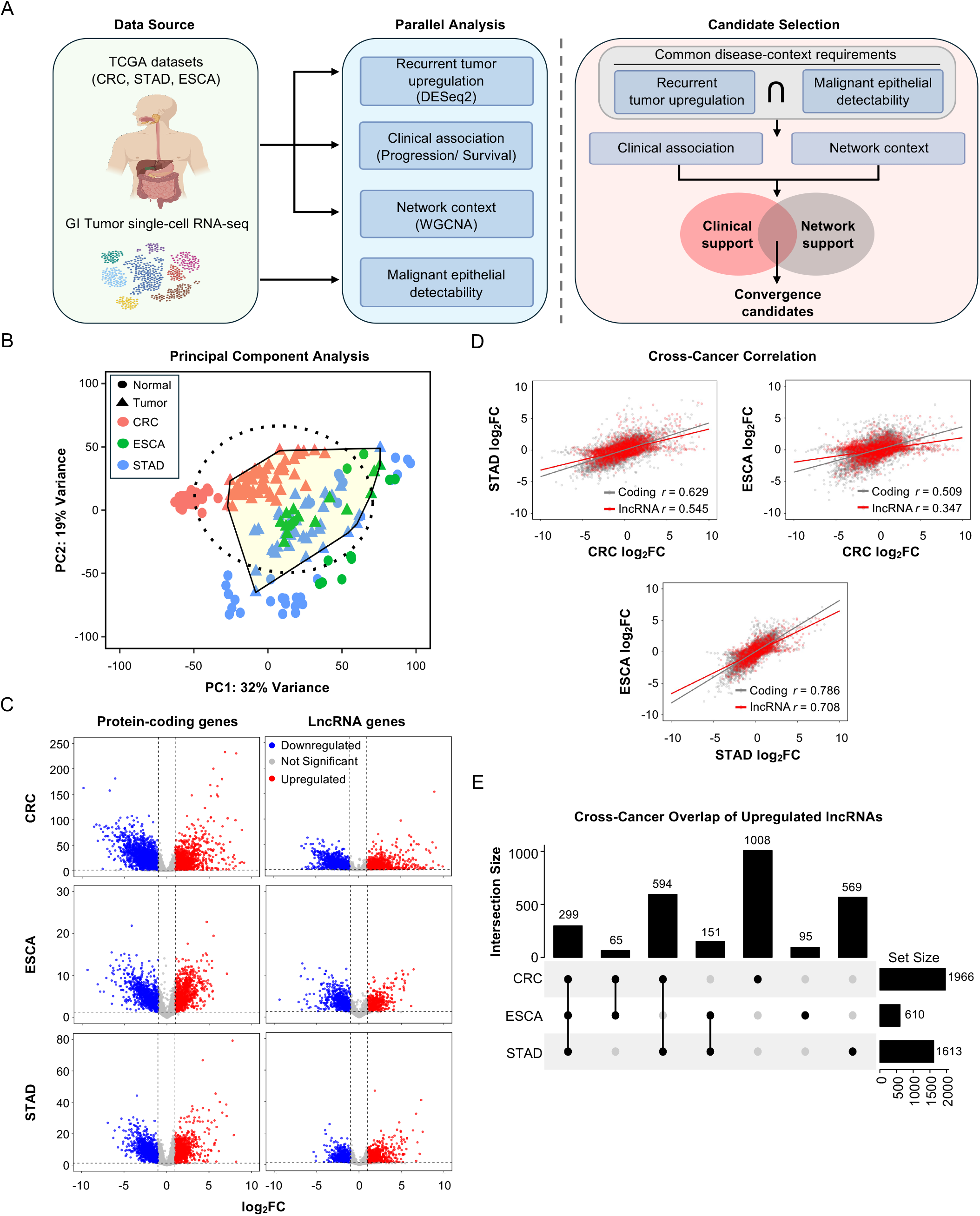
Pan-GI cancers exhibit lncRNA dysregulation. (A) Overview of the pan-GI lncRNA prioritization strategy. TCGA tumor–normal expression profiles and GI tumor single-cell RNA-seq data from CRC, STAD, and ESCA were evaluated separately across four types of evidence: recurrent tumor upregulation, clinical association with disease progression or survival, network context from weighted gene co-expression network analysis (WGCNA), and malignant epithelial detectability. Recurrent tumor upregulation and malignant epithelial detectability served as common disease-context requirements, whereas clinical association and network context provided parallel forms of additional support. Candidates supported by both the clinical and network axes were designated convergence candidates. (B) PCA of variance-stabilized TCGA expression profiles from CRC, STAD, and ESCA tumor and matched normal samples. Dashed contour and shaded polygon mark the tumor sample distribution. (C) Paired tumor–normal differential expression of protein-coding genes and lncRNAs in CRC, STAD, and ESCA. Differential expression was analyzed using DESeq2 with apeglm-shrunken log2 fold-change (log2FC) estimates. Red, blue, and grey points indicate significantly upregulated, significantly downregulated, and non-significant genes, respectively. Significance was defined as absolute shrunken log2FC ≥ 0.5 and false discovery rate (FDR) < 0.05. (D) Pairwise cross-cancer comparison of tumor–normal shrunken log2FC estimates. Pearson correlations were calculated separately for protein-coding genes and lncRNAs using genes tested in both cohorts; lines indicate linear regression fits. (E) UpSet analysis of tumor-upregulated lncRNAs across CRC, STAD, and ESCA, defined by shrunken log2FC ≥ 0.5 and FDR < 0.05. Bar plots show cohort-specific and intersecting lncRNA sets, including 299 lncRNAs upregulated in all three cohorts and 1,109 lncRNAs upregulated in at least two cohorts. Related cohort information, differential-expression results, cross-cancer correlations, and recurrent lncRNA sets are provided in Table S1.

To define the tumor-associated transcriptional landscape, we performed paired tumor–normal differential expression analyses within each cancer type (|shrunken log2FC| ≥ 0.5, FDR < 0.05). Both protein-coding and lncRNA genes showed widespread dysregulation across CRC, STAD, and ESCA (Fig. 1C; Tables S1B and S1C). Among lncRNAs, 1,966 were upregulated and 1,500 downregulated in CRC, 1,613 up and 990 down in STAD, and 610 up and 945 down in ESCA. Representative proliferation, metabolic, and ribosome/translation-associated markers showed expected tumor-associated patterns, with housekeeping genes included as a reference class (Fig. S1B). Whereas protein-coding changes were dominated by downregulation in all three cancers, lncRNA changes were skewed toward upregulation in CRC and STAD but toward downregulation in ESCA, defining a broad tumor-associated lncRNA landscape with substantial cancer-context dependence.

To determine how consistently these tumor-associated changes were shared across GI cancers, we compared apeglm-shrunken tumor–normal fold changes across cancer types. Protein-coding genes exhibited moderate-to-strong concordance (Pearson *r* = 0.51–0.79), whereas lncRNA concordance was lower (*r* = 0.35–0.71; Fig. 1D; Table S1D). Concordance for both gene classes was highest between STAD and ESCA and lowest between CRC and ESCA. The lower lncRNA concordance was consistent with greater tissue- and context-specificity of the lncRNA transcriptome (14, 15), indicating that recurrent pan-GI lncRNA signals are embedded within a strongly cancer-dependent transcriptional background.

Given the lower cross-cancer concordance, we focused on recurrent tumor-upregulated lncRNAs to identify candidates with potential tumor-promoting activity. Cross-cohort intersection identified 299 lncRNAs upregulated in all three cancers and 1,109 upregulated in at least two of three cohorts (Fig. 1E; Tables S1E and S1F), representing approximately 11% and 40%, respectively, of the union of tumor-upregulated lncRNAs. Thus, cross-cancer recurrence substantially restricted the broader tumor-associated lncRNA landscape and defined the recurrent tumor-upregulation set carried forward for multidimensional prioritization.

### Identification of lncRNAs associated with adverse patient outcomes

To determine how tumor-associated lncRNA expression relates to disease progression and patient outcome, we evaluated clinicopathologic stage and overall survival across GI cohorts (Fig. 2A; Tables S2A and S2B). For stage analysis, DESeq2 likelihood ratio testing (LRT) identified lncRNAs whose expression varied across primary tumor extension (T), regional nodal involvement (N), or distant metastasis (M) within each cancer type (Fig. 2B; Table S2C). Cross-cohort recurrence was greatest for T stage, with 2,476 lncRNAs supported by significant LRT in at least two cohorts and 430 in all three, followed by N stage (1,439 and 172) and M stage (476 and 22). Thus, primary tumor extension yielded the most recurrent stage-associated lncRNA signals across GI cancers, whereas M-stage associations were more restricted.

**Figure 2.**
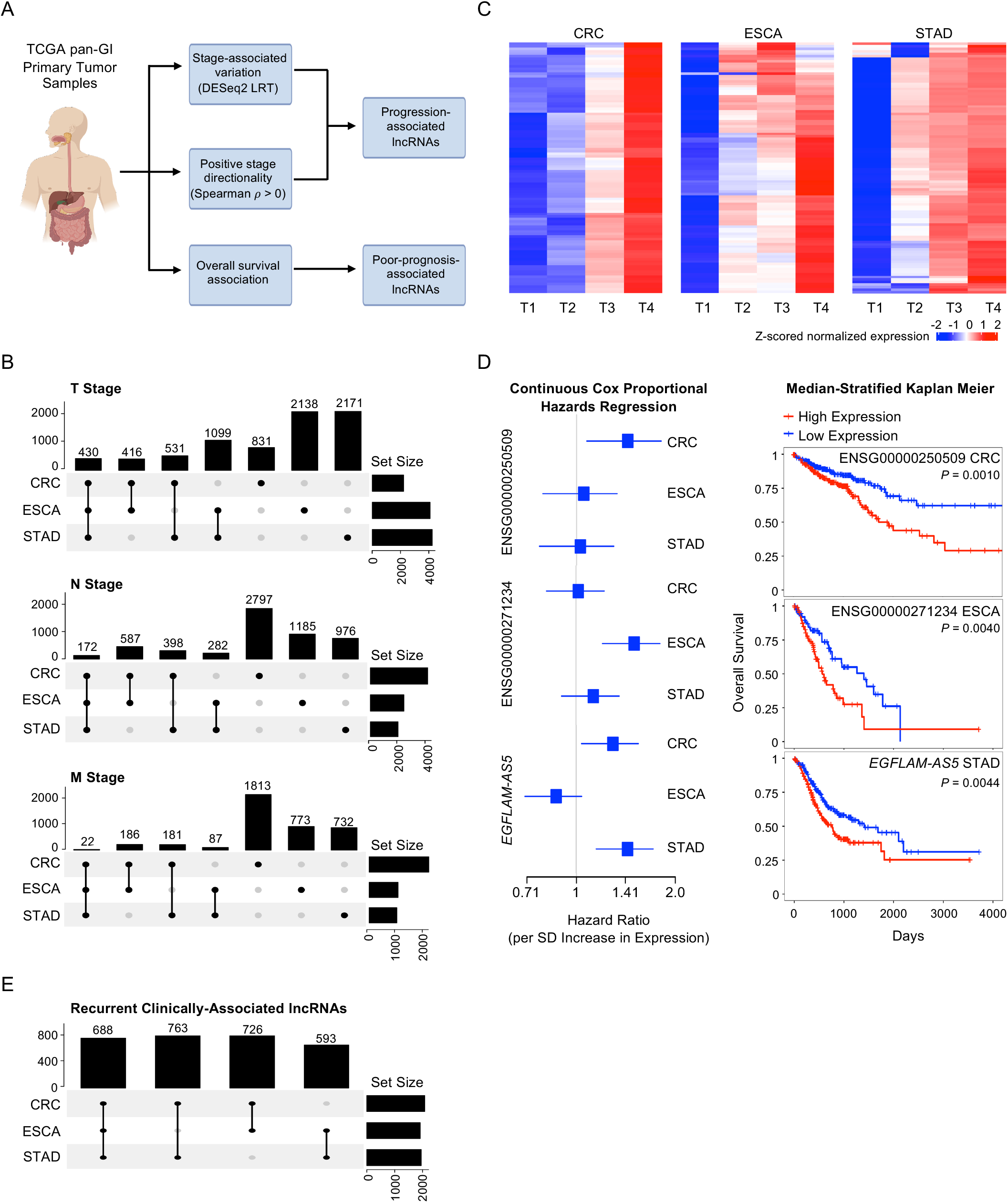
Identification of lncRNAs associated with adverse patient outcomes. (A) Overview of the TCGA-based clinical prioritization strategy. Primary tumor RNA-seq datasets from CRC, STAD, and ESCA were analyzed to identify lncRNAs associated with AJCC T, N, and M stages and overall survival. (B) DESeq2 LRT identified lncRNAs whose expression varied across T, N, or M stage categories within each cancer cohort. UpSet plots summarize recurrent and cohort-specific stage-associated lncRNAs. (C) Heatmaps showing representative T stage-ordered expression patterns of stage-associated lncRNAs in CRC, ESCA, and STAD. Samples are ordered by AJCC T stage, and expression values are shown after normalization and row scaling. Genes shown were filtered by LRT *P* < 0.05 and ranked by positive Spearman correlation with stage. (D) Survival association of representative lncRNAs across GI cancer cohorts. Forest plots show hazard ratios and confidence intervals from univariate Cox proportional hazards regression, with hazard ratios representing the change in risk per one standard deviation increase in normalized expression. Kaplan-Meier curves show overall survival after stratification into high- and low-expression groups using cohort-specific median expression. (E) Integrated UpSet analysis of clinically prioritized lncRNAs. Recurrent clinically prioritized candidates were defined as lncRNAs supported by progression evidence in at least two GI cancer cohorts and/or poor-survival evidence in at least two GI cancer cohorts. The analysis identified 2,770 recurrent clinically prioritized candidates, including a 688-lncRNA core supported across CRC, STAD, and ESCA. Related sample counts and clinical analysis results are provided in Table S2.

Because LRT identifies stage-associated variation without specifying direction, we used Spearman correlation to distinguish lncRNAs whose expression increased with advancing disease (Fig. S2A; Table S2C). Progression-linked lncRNAs were defined by LRT-supported stage variation together with positive stage directionality, with Spearman correlation used to assign direction rather than as an additional significance filter. Stage-ordered heatmaps showed increasing expression of representative T- and N-stage candidates while retaining cohort-specific patterns (Figs. 2C and S2B); M-stage heatmaps were not generated because M stage was binary (M0/M1). Across cohorts, recurrence declined from T to N to M stage: 1,252 of 5,160 T-stage candidates (24.3%) were supported in at least two cohorts and 243 in all three, compared with 556 of 3,889 N-stage candidates (14.3%; 37 in all three) and 78 of 1,615 M-stage candidates (4.8%; two in all three) (Fig. S2C; Table S2C). Limited M-stage recurrence occurred in the context of smaller M1 groups, particularly in STAD and ESCA (Table S2B). These results identified progression-linked lncRNAs while underscoring their strong cancer-context dependence.

To complement progression-associated evidence with patient outcome, we evaluated lncRNA expression in relation to overall survival. Univariate Cox proportional hazards analysis identified 831 lncRNAs associated with poor survival in CRC, 2,466 in STAD, and 387 in ESCA (Table S2D). Forest plots and Kaplan-Meier curves of representative candidates, ENSG00000250509 in CRC, *EGFLAM-AS5* in STAD, and ENSG00000271234 in ESCA, showed reduced overall survival with high expression (Fig. 2D; Table S2D). We then integrated directional progression and poor-survival evidence to define recurrent clinically prioritized lncRNAs. Candidates were retained if they showed progression support in at least two GI cancer cohorts or poor-survival support in at least two cohorts (Table S2E). This identified 2,770 recurrent clinical candidates, including 688 supported across all three cohorts (24.8% of the recurrent set; Fig. 2E; Table S2E). Previously implicated GI cancer lncRNAs, including *IGF2-AS* and *PART1* (21, 22), were recovered, supporting the analysis strategy (Table S2E). Altogether, these analyses resolved widespread cohort-level clinical associations into a recurrent set of lncRNAs linked to adverse disease features across GI cancers.

### Co-expression network analysis places lncRNAs within disease-associated transcriptional modules

To place lncRNAs within coordinated transcriptional programs, we performed weighted gene co-expression network analysis (WGCNA) on pooled TCGA primary tumors from CRC, STAD, and ESCA (Figs. 3A and S3A–C). Hierarchical clustering identified 21 non-grey modules whose eigengenes showed distinct associations with cancer type and tumor features (Fig. 3B; Table S3A). We evaluated pooled modules for disease-context association, lncRNA representation, and lncRNA connectivity. Sixteen non-grey modules showed at least one FDR-significant module-trait association and contained at least one lncRNA with eigengene-based connectivity (kME) ≥ 0.5 (Fig. 3C; Table S3A). Functional annotation of these modules identified programs related to extracellular matrix (ECM) organization, cytoplasmic translation, RNP complex biogenesis, immune signaling, chromosome segregation, endosomal transport, epidermis development, and antiviral response (Table S3B).

**Figure 3.**
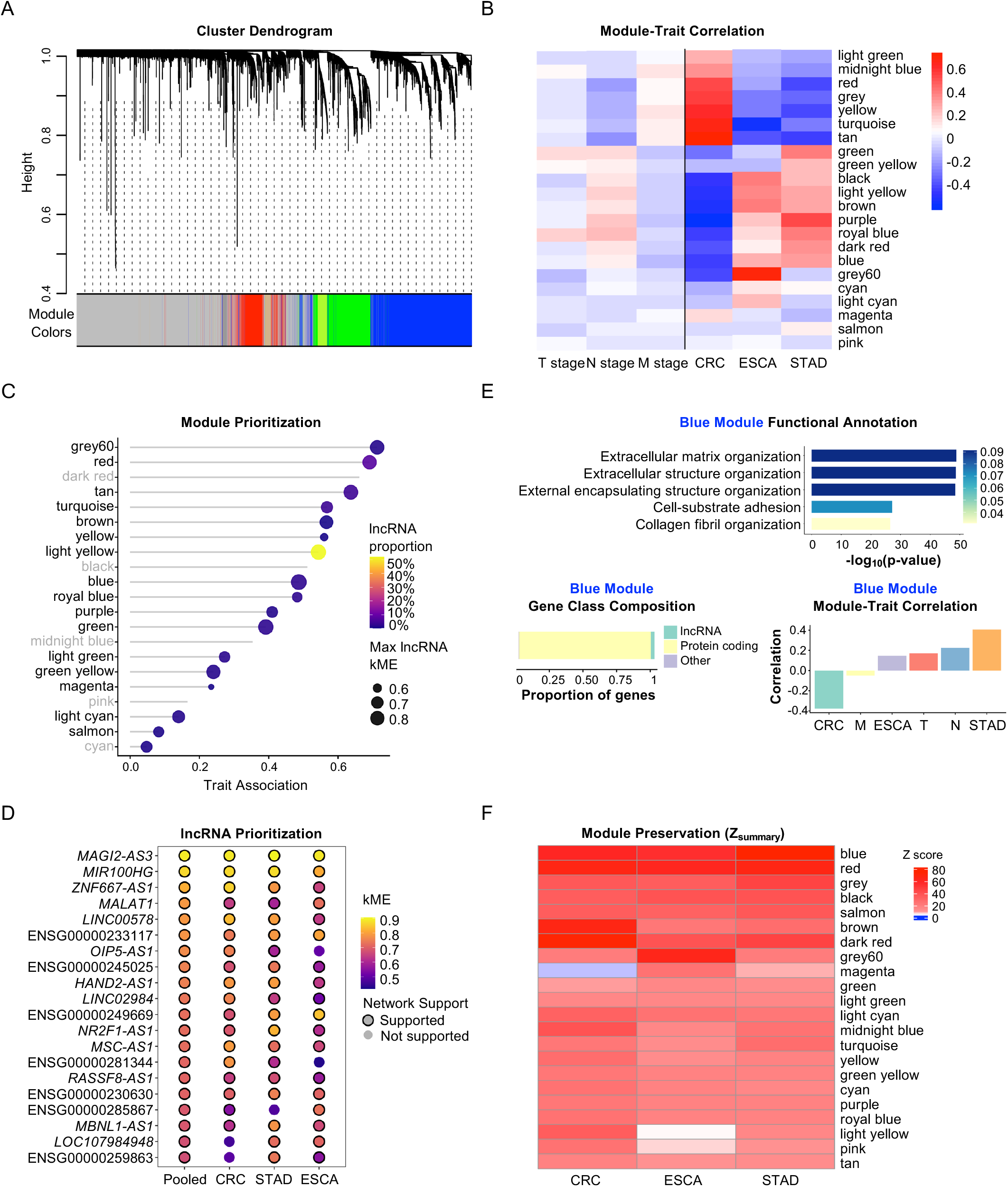
Co-expression network analysis places lncRNAs within disease-associated transcriptional modules. (A) WGCNA of pooled TCGA primary tumor RNA-seq data from CRC, STAD, and ESCA. Gene dendrogram shows hierarchical clustering based on co-expression topology, with module assignments indicated by color. (B) Module–trait correlation heatmap showing associations between WGCNA module eigengenes and cohort or tumor-feature variables. Color intensity represents Pearson correlation coefficients. (C) Network features of pooled WGCNA modules. Modules are shown according to module-trait association, lncRNA representation, and maximum lncRNA kME. Prioritized modules had at least one FDR-significant module-trait association and at least one lncRNA with kME ≥ 0.5. Trait association represents the maximum absolute module–trait correlation across cohort and tumor-feature variables. Prioritized modules are labeled in black and non-prioritized modules in grey. (D) Top 20 lncRNAs assigned to network-prioritized pooled pan-GI WGCNA modules, ranked by absolute pooled kME. Dot color indicates kME, and cancer-type-resolved support across CRC, STAD, and ESCA is shown as a network robustness annotation. Related module annotations, functional enrichment results, and lncRNA rankings are provided in Table S3. (E) Functional annotation of the prioritized blue module. Bar plot shows representative GO-BP terms enriched among blue-module genes. Lower panels show gene-class composition and module–trait correlation patterns for the blue module. (F) Cross-cohort preservation analysis of pooled pan-GI WGCNA modules in CRC, STAD, and ESCA. Heatmap shows module-preservation Zsummary scores across the three cancer cohorts.

To identify which individual lncRNAs were most strongly supported by network analysis, we ranked the 216 lncRNAs within prioritized modules by kME and assessed their support in cancer-type-resolved networks (Fig. 3D; Table S3C). Of these, 162 showed network support in at least two GI cancer cohorts and 88 in all three. High-ranking candidates were concentrated in the blue module, which contained 16 of the top 20 lncRNAs, and 15 of the top 20 were supported across all three cohorts. The highest-ranked candidates included established cancer-associated lncRNAs such as *MAGI2-AS3*, *MIR100HG*, and *MALAT1* (23–25), supporting the ability of the analysis to recover disease-relevant noncoding transcripts.

Because the blue module contained most of the highest-ranked recurrent candidates, we examined its biological composition as a representative example. Blue-module genes were enriched for ECM organization, extracellular structure organization, cell-substrate adhesion, and collagen-associated processes (Fig. 3E). Cancer-type-resolved networks showed cohort-specific module-trait relationships (Figs. S3D–F), whereas preservation analysis indicated that the pooled modules were largely reproducible across individual cancer types (Fig. 3F; Table S3A). Seventeen of 21 pooled modules showed strong preservation across all three cohorts, including 13 of 16 prioritized modules. Thus, network analysis identified recurrently supported lncRNAs within biologically interpretable transcriptional programs that were largely preserved across GI cancers.

### Single-cell RNA-seq analysis identifies malignant epithelial-associated lncRNA candidates in GI cancer

Bulk tumor profiles cannot distinguish lncRNA expression arising from malignant epithelial cells from that contributed by stromal, immune, endothelial, and other cell populations. We therefore analyzed CRC, STAD, and ESCA single-cell profiles from the TabulaTIME pan-cancer single-cell atlas (26) to resolve candidate expression at the cell-type level. The GI dataset comprised 14,903 single-cell-derived metacell profiles from 237 patient samples, including 2,648 malignant epithelial profiles, with major cell-type assignments supported by canonical lineage markers (Figs. 4A, S4A, and S4B; Table S4A).

**Figure 4.**
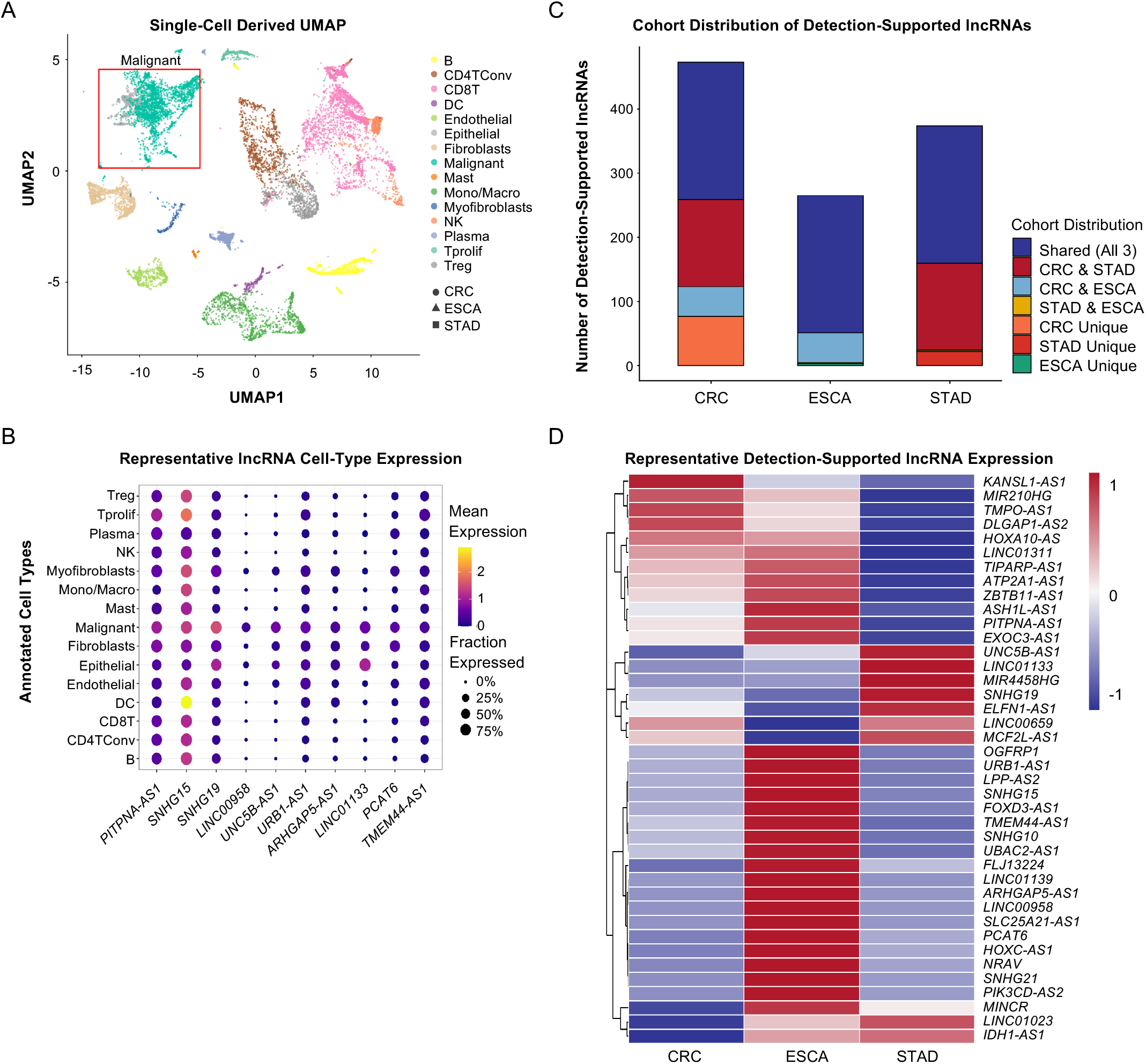
Single-cell RNA-seq analysis identifies malignant epithelial-associated lncRNA candidates in GI cancer. (A) Integrated UMAP of single-cell-derived GI cancer profiles from the TabulaTIME pan-cancer atlas, including CRC, STAD, and ESCA. Profiles are colored by major cell-type annotation and shaped by cancer cohort. The red box indicates the malignant epithelial compartment used for downstream lncRNA prioritization. (B) Dot plot showing expression of representative lncRNA candidates across major annotated cell types. Dot size indicates the fraction of profiles expressing each lncRNA, and color indicates average normalized expression. The candidate lncRNAs shown represent top-ranked malignant-associated candidates from the single-cell analysis. (C) Detection-based summary of lncRNA expression across CRC, STAD, and ESCA. LncRNAs detected in at least 1% of malignant profiles were classified as detection-supported. (D) Clustered heatmap showing row-scaled average normalized expression of representative detection-supported lncRNA candidates across CRC, STAD, and ESCA. Related dataset composition, annotation metadata, and candidate-level single-cell prioritization metrics are provided in Table S4.

To distinguish lncRNAs preferentially associated with malignant epithelial cells from those more broadly expressed across the tumor microenvironment, we performed patient-level pseudobulk malignant-versus-other differential expression analysis within each cancer type. This identified 84 unique malignant-associated lncRNAs with significant differential expression in at least one cancer type (Table S4B). Differential-expression support was strongly skewed toward ESCA (CRC, n = 4; STAD, n = 6; ESCA, n = 79), revealing substantial context dependence across GI cancer types. Consistent with this pattern, only five lncRNAs, *ELFN1-AS1*, *FAM201A*, *FOXD3-AS1*, *LINC00853*, and *RHPN1-AS1*, showed malignant-associated differential expression in two of the three GI cancer contexts, and none did so across all three. To examine the cell-type distribution of these malignant-associated signals, we profiled representative high-ranking candidates across major annotated cell populations. These candidates were preferentially expressed in malignant epithelial populations (Fig. 4B). Therefore, malignant-versus-other differential expression identified lncRNAs with preferential malignant epithelial expression, but these signals showed limited recurrence across GI cancer contexts.

Differential expression reflects preferential enrichment rather than transcript presence, and sparse capture can obscure low-abundance lncRNAs in single-cell data (6, 7, 27, 28). We therefore assessed malignant epithelial detectability separately, defining detection support as expression in at least 1% of malignant profiles within a cancer context. This identified 500 unique lncRNAs, including 398 detected in at least two GI cancer contexts and 214 across all three (Fig. 4C; Table S4B). We next evaluated detectability of the 84 unique malignant-associated lncRNAs across CRC, STAD, and ESCA, yielding 252 lncRNA-cancer contexts. All 89 contexts with significant differential expression support met the detection threshold, while 141 of the remaining 163 non-significant contexts were also detection-supported, such that 230 of 252 (91.3%) lncRNA-cancer contexts showed malignant epithelial detectability, indicating that lack of significant malignant enrichment generally did not reflect absence from the malignant compartment (Table S4B). Clustering of representative detection-supported lncRNAs across CRC, STAD, and ESCA revealed both broadly shared and cancer-type-enriched expression patterns (Fig. 4D), indicating that broad malignant epithelial detectability did not imply uniform expression across GI cancers. Collectively, malignant-versus-other differential enrichment was highly context-dependent, whereas malignant epithelial detectability extended across a substantially broader range of GI cancer contexts.

### Multidimensional prioritization identifies *LINC01094* as a nuclear-enriched lncRNA associated with a distinct CRC transcriptional state

Having defined tumor-associated expression, clinical association, co-expression network context, and malignant epithelial detectability separately, we integrated these analyses only at candidate selection (Fig. 1A). Recurrent tumor upregulation in at least two cohorts together with malignant epithelial detectability in at least two cancer contexts identified 69 lncRNAs that met the common disease-context requirements (Table S5A). Among these, 24 had clinical support and 12 had network support (Fig. 5A). Five candidates, *MIAT*, *ITGB2-AS1*, *HOXA11-AS*, *LINC01094*, and *LINC01235*, were supported by both analyses. This convergence reduced the broader disease-associated set to a small group of candidates supported across complementary evidence dimensions.

**Figure 5.**
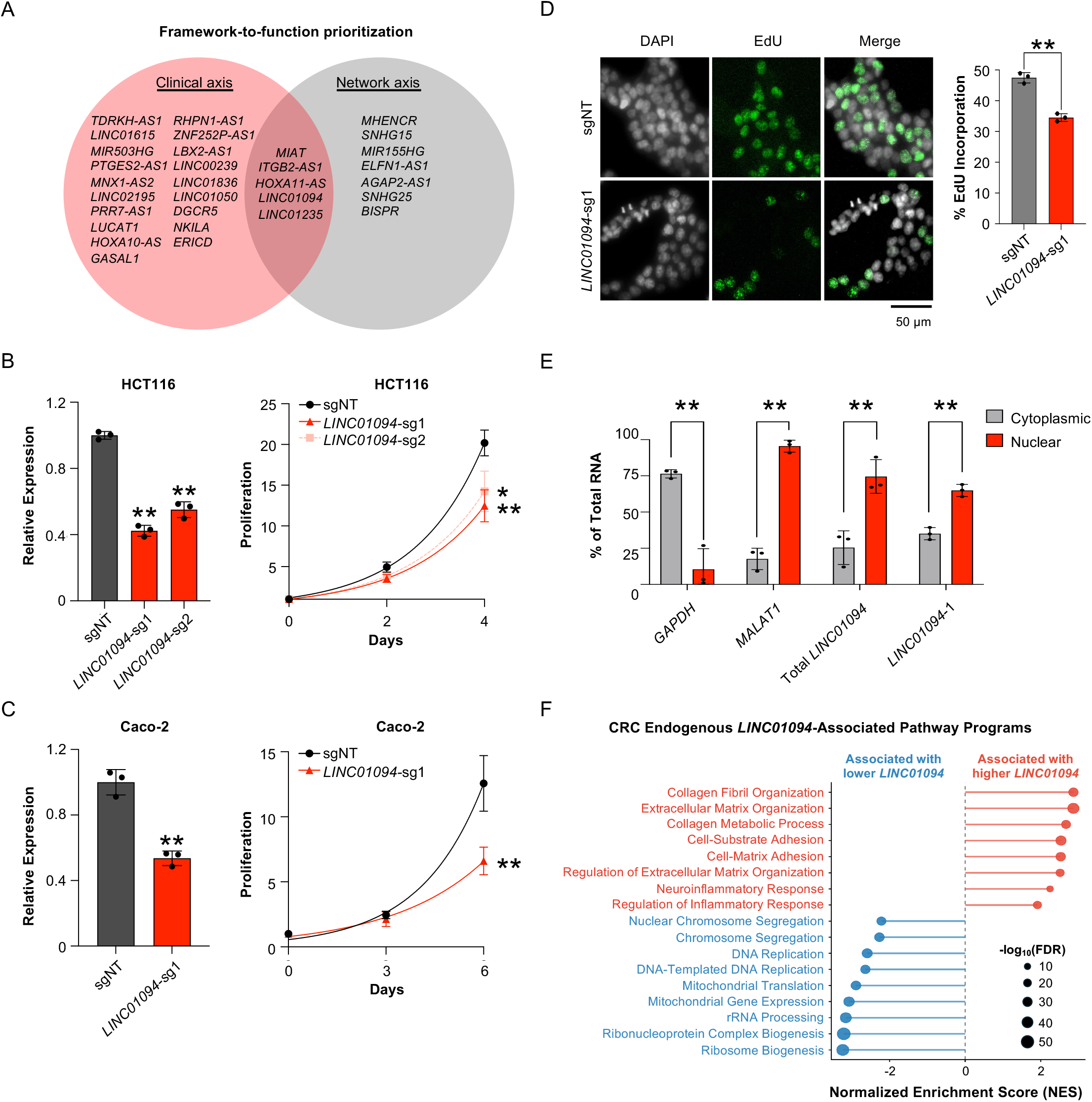
Multidimensional prioritization identifies *LINC01094*, a nuclear-enriched lncRNA linked to a colorectal cancer transcriptional state. (A) Candidate-selection strategy. Both axes required recurrent tumor upregulation (shrunken log2FC ≥ 0.5, FDR < 0.05 in ≥ 2 of 3 cohorts) and malignant epithelial detectability (≥1% of malignant profiles in ≥ 2 cancer contexts). Clinical support required progression or poor-survival in ≥ 2 cohorts, whereas network support required assignment to a network-prioritized WGCNA module. These criteria defined parallel candidate sets, with five candidates shared by both. (B) RT-qPCR validation of CRISPRi-mediated *LINC01094* depletion (left) and growth analysis (right) of HCT116 cells expressing dCas9-KRAB and either a non-targeting control sgRNA (sgNT) or one of two independent *LINC01094*-targeting sgRNAs. *LINC01094* abundance was reduced by approximately 60% and 45% with *LINC01094-*sg1 and *LINC01094*-sg2, respectively. Because *LINC01094-*sg1 produced greater knockdown, it was used for selected follow-up assays. Cell growth was measured over four days following selection. Data are presented as mean ± SD (n=3 biological replicates). \**P* <0.05; \*\**P* <0.01, calculated by two-tailed t-test. (C) RT-qPCR validation of CRISPRi-mediated *LINC01094* depletion (left) and growth analysis (right) of Caco-2 cells expressing dCas9-KRAB and either sgNT or *LINC01094-*sg1. *LINC01094* abundance was reduced by approximately 50% with *LINC01094-*sg1. Cell growth was measured over six days following selection. Data are presented as mean ± SD (n=3 biological replicates). \**P* <0.05; \*\**P* <0.01, calculated by two-tailed t-test. (D) Fluorescence microscopy (left) and quantification (right) of DAPI and EdU incorporation following *LINC01094* depletion with *LINC01094-*sg1 in HCT116 cells. Data are presented as mean ± SD (n=3 biological replicates). \**P* <0.05; \*\**P* <0.01, calculated by two-tailed t-test. Scale bar shows 50 μm. (E) Subcellular fractionation followed by RT-qPCR analysis of *LINC01094* localization in HCT116 cells. *LINC01094* was enriched in the nuclear fraction using an isoform-shared primer set and a *LINC01094*-1 specific primer set. *MALAT1* and *GAPDH* served as nuclear and cytoplasmic controls, respectively. Data are presented as mean ± SD (n=3 biological replicates). \**P* <0.05; \*\**P* <0.01, calculated by two-tailed t-test. (F) CRC-restricted CCLE continuous-expression GSEA across 52 CRC cell lines. Genes were ranked according to their association with endogenous *LINC01094* expression, and representative high-NES GO-BP terms were selected from FDR-significant pathways within major positive- and negative-NES pathway classes. Positive NES indicates pathways associated with higher endogenous *LINC01094* expression, whereas negative NES indicates pathways associated with lower endogenous *LINC01094* expression. Dot size indicates enrichment significance as shown.

We focused on *LINC01094* for functional characterization because its intergenic location and minimal overlap with neighboring transcripts facilitated downstream experimental analysis. Prior studies have linked *LINC01094* to CRC progression, proliferation and invasion (20). Consistent with this, analysis of TCGA data further confirmed that *LINC01094* is elevated in CRC tumors relative to matched normal tissues and increases with advancing T stage (Figs. S5A and S5B). CRISPRi-mediated depletion of *LINC01094* significantly impaired cell growth in both HCT116 and Caco-2 cells (Figs. 5B and 5C). 5-ethynyl-2′-deoxyuridine (EdU) incorporation was also reduced in *LINC01094*-deficient HCT116 cells, indicating reduced DNA synthesis (Fig. 5D). Together, these findings show that *LINC01094* supports cell growth across two CRC models.

Because lncRNA loci frequently generate multiple transcript products, we next defined the *LINC01094* transcript expressed in HCT116 cells. Public long-read RNA-seq datasets (29) integrated with ENCODE4 long-read transcript annotations (30) supported two alternatively spliced candidate *LINC01094* isoforms (Fig. S5C; Table S5B). RT-PCR and isoform-directed RT-qPCR showed that the predominant CRISPRi-sensitive product was consistent with *LINC01094*-1, whereas the predicted *LINC01094*-2 transcript was present at substantially lower abundance (Figs. S5D and S5E). Subcellular fractionation further demonstrated nuclear enrichment using both isoform-shared and *LINC01094*-1-specific primer sets (Fig. 5E).

Although *LINC01094* has been linked to CRC progression and proliferation, the transcriptional state associated with its endogenous expression in CRC remained incompletely defined. We therefore analyzed 52 CRC cell lines from the Cancer Cell Line Encyclopedia (CCLE). HCT116 fell within the observed *LINC01094* expression distribution (Fig. S5F; Table S7A). Continuous-expression gene set enrichment analysis (GSEA) revealed that higher *LINC01094* expression was associated with ECM/collagen organization, cell-matrix and cell-substrate adhesion, and inflammatory-response pathways, whereas lower expression was associated with ribosome/RNP biogenesis, rRNA processing, mitochondrial gene expression and translation, DNA replication, and chromosome segregation (Fig. 5F). Altogether, these findings define a coherent endogenous *LINC01094*-associated transcriptional axis in CRC, with higher and lower *LINC01094* expression marking opposing ECM/adhesion-inflammatory and replication-mitochondrial-ribosome-associated programs, respectively.

### Acute *LINC01094* depletion opposes the endogenous CRC transcriptional state and induces replication-associated checkpoint activation

To test whether acute *LINC01094* loss perturbs the endogenous CRC transcriptional state, we performed RNA-seq following CRISPRi with two independent sgRNAs, each compared with non-targeting control cells (Table S6A). PCA clearly separated *LINC01094*-depleted cells from control cells (PC1, 78.7% variance; Fig. S6A). Across 14,724 detected genes, sgRNA responses were strongly correlated (Pearson *r* = 0.83; Fig. 6A); of 1,550 genes significantly altered by both sgRNAs, 1,544 (99.6%) changed in the same direction, with a Pearson correlation of 0.95 within this shared set (Tables S6B-S6D). GSEA identified 532 positively and 681 negatively enriched FDR-significant GO-BP terms (Table S6E). Induced programs included chromosome segregation, DNA replication, mitochondrial gene expression and translation, and ribosome/RNP biogenesis, whereas adhesion, ECM/collagen, and inflammatory/MHC-associated programs were suppressed (Fig. 6B). RT-qPCR analysis of 15 representative genes confirmed induction of replication/checkpoint- and DNA damage response (DDR)-associated genes and suppression of ECM/collagen- and inflammatory/MHC-associated genes in HCT116 cells (Fig. 6C). In Caco-2 cells, seven HCT116-responsive markers showed concordant directional changes, with *TGM2* and *FN1* significantly reduced (Fig. S6B). *SLPI*, a reported effector of the cytoplasmic miR-1266-5p/SLPI ceRNA model in CRC (20), showed only a modest, non-significant decrease in HCT116 RNA-seq after either sgRNA and no detectable change by RT-qPCR in either HCT116 or Caco-2 cells (Fig. S6C; Table S6B). Thus, *LINC01094* depletion drives coordinated remodeling of opposing transcriptional programs in CRC cells without a prominent *SLPI*-associated response.

**Figure 6.**
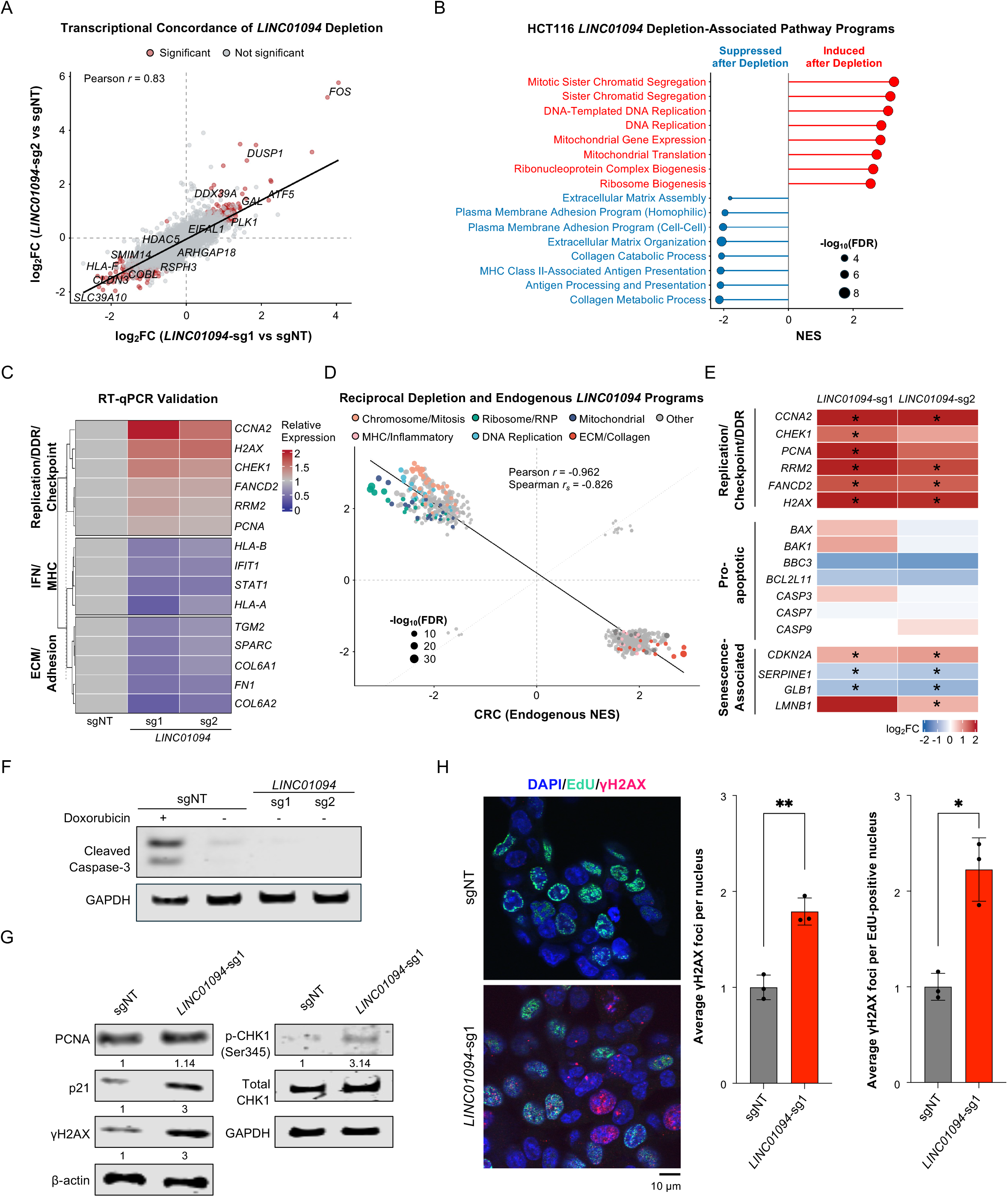
Acute *LINC01094* depletion opposes the endogenous CRC transcriptional state and induces replication-associated checkpoint activation (A) Transcriptome-wide concordance of the responses to two independent *LINC01094*-targeting sgRNAs. Scatter plot compares gene-level log2FC values for each *LINC01094*-targeting sgRNA relative to sgNT across 14,724 detected genes. Pearson r = 0.83. Genes meeting the indicated differential-expression criteria in both sgRNA comparisons are highlighted, and selected highly concordant genes are labeled. (B) GSEA of the acute HCT116 *LINC01094*-depletion response. Genes were ranked by the differential expression response to *LINC01094* depletion, and representative high-NES GO-BP terms were selected from FDR-significant pathways (FDR <0.05) within major positive- and negative-NES pathway classes. Positive NES indicates pathways induced after *LINC01094* depletion, whereas negative NES indicates pathways suppressed after *LINC01094* depletion. Dot size indicates enrichment significance as shown; full FDR-significant GSEA results are provided in Table S6E. (C) RT-qPCR validation of representative RNA-seq marker genes following *LINC01094* depletion. Heatmap shows mean relative expression normalized to sgNT cells from three biological replicates. Replication/checkpoint and DDR-associated markers (*CCNA2*, *CHEK1*, *PCNA*, *RRM2*, *FANCD2*, and *H2AX*) were induced, whereas ECM/collagen-associated markers (*SPARC*, *TGM2*, *COL6A1*, *COL6A2*, and *FN1*) and inflammatory/MHC-associated markers (*HLA-A*, *HLA-B*, *STAT1*, and *IFIT1*) were suppressed. (D) Direct comparison of CRC CCLE endogenous *LINC01094*-associated programs with the acute HCT116 *LINC01094*-depletion response. Scatter plot shows NES values for GO terms significantly enriched in both analyses. Among 764 GO terms significant in both analyses, 748 (97.9%) showed opposite directionality (Pearson *r* = −0.962; Spearman *r*s = -0.826). (E) Heatmap of select replication/checkpoint/DDR-, apoptosis-, and senescence-associated regulators in the HCT116 *LINC01094*-depletion RNA-seq dataset. Values show gene-level log2FC for each *LINC01094*-targeting sgRNA relative to sgNT. Asterisks indicate FDR < 0.05 in the corresponding comparison. (F) Western blot analysis of cleaved caspase-3 in HCT116 cells expressing sgNT or either *LINC01094*-targeting sgRNA. Doxorubicin-treated sgNT served as a positive control for caspase-3 cleavage, and GAPDH served as the loading control. (G) Western blot analysis of PCNA, p21, γH2AX, p-CHK1 Ser345 and total CHK1 following *LINC01094* depletion with *LINC01094*-sg1 in HCT116 cells. β-actin and GAPDH served as the loading controls as indicated. Densitometric values were normalized to the corresponding loading or total-protein controls and scaled to sgNT. Blots in panels F and G were cropped for presentation; uncropped original membrane sections are provided in the Supplementary Information. (H) Immunofluorescence analysis of DAPI, EdU, and γH2AX following *LINC01094* depletion with *LINC01094*-sg1 in HCT116 cells. Quantification shows average γH2AX foci per nucleus and average γH2AX foci per EdU-positive nucleus. Data are presented as mean ± SD (n=3 biological replicates). \**P* <0.05; \*\**P* <0.01, calculated by two-tailed t-test. Scale bar shows 10 μm.

We next compared the acute HCT116 CRISPRi response with the independently derived endogenous *LINC01094*-associated state from cross-line variation across 52 CRC models. Across the 764 GO terms significant in both analyses, normalized enrichment scores (NES) were inversely correlated (Pearson *r* = −0.962; Spearman *rs* = −0.826; Fig. 6D), with 748 (97.9%) showing opposite directionality (Table S7B). Among the major pathway classes represented in both analyses, ECM/adhesion and inflammatory pathways were associated with higher endogenous *LINC01094* expression but were suppressed following depletion, whereas DNA replication, chromosome segregation, mitochondrial, and ribosome/RNP programs showed the opposite pattern. Thus, acute *LINC01094* depletion largely opposes the endogenous *LINC01094*-associated transcriptional state across CRC models.

We further assessed the robustness and lineage specificity of this reciprocal relationship using alternative expression stratifications in CRC and corresponding analyses in STAD and ESCA. Threshold- and median-stratified CRC analyses preserved the same inverse pathway relationship (Fig. S6D; Table S7C). In contrast, applying the same framework to other GI lineages did not reproduce the full CRC-associated architecture. STAD partially recapitulated the CRC ECM/adhesion and immune-associated programs, whereas ESCA showed a distinct profile characterized by cell-cycle/checkpoint and keratinization/squamous-associated terms (Fig. S6E; Tables S7D and S7E). Therefore, the reciprocal relationship is reproducible within CRC and robust to expression stratification, but is not fully recapitulated across other GI lineages, supporting a context-dependent relationship between *LINC01094* expression and transcriptional state.

A major component of the reciprocal state shift was induction of replication/checkpoint and DDR programs. However, this transcriptional response occurred despite reduced cell growth and DNA synthesis, suggesting that the shift toward replication-associated programs did not reflect productive cell-cycle activation. We therefore examined replication/checkpoint-, apoptosis-, and senescence-associated regulators across both *LINC01094*-targeting RNA-seq datasets (Fig. 6E; Tables S6B and S6C). Replication/checkpoint- and DDR-associated genes were coordinately induced, whereas apoptosis-associated regulators showed no coordinated transcriptional response. Senescence-associated markers were also discordant: *CDKN2A* increased with both sgRNAs, *SERPINE1* and *GLB1* decreased with both, and *LMNB1* increased with sg1 but was unchanged with sg2. Consistent with the absence of a coordinated apoptotic response, cleaved caspase-3 was not detected following *LINC01094* depletion with either sgRNA, while doxorubicin-treated control cells showed robust caspase-3 cleavage (Fig. 6F). These findings are consistent with checkpoint activation rather than a coordinated apoptotic or senescence response.

Next, we examined DNA damage and checkpoint signaling following *LINC01094* depletion. PCNA increased modestly, accompanied by increased p21 and γH2AX (Fig. 6G), consistent with activation of a DNA damage and checkpoint response. Notably, p21 protein accumulation occurred without a corresponding increase in *CDKN1A* mRNA (Fig. 6G; Tables S6B and S6C). *LINC01094* depletion also increased CHK1 Ser345 phosphorylation relative to total CHK1, further supporting checkpoint activation. Because *LINC01094* depletion reduced DNA synthesis, we combined EdU labeling with γH2AX immunofluorescence to determine whether DNA-damage signaling was also evident in cells that remained engaged in DNA synthesis. γH2AX foci increased by approximately 1.8-fold across all nuclei and by approximately twofold within residual EdU-positive nuclei (Fig. 6H), with doxorubicin-treated cells serving as a positive control for γH2AX induction (Fig. S6F). Thus, DNA-damage signaling remained elevated even in cells that continued to incorporate EdU. Together, these findings show that acute *LINC01094* loss uncouples induction of replication-associated transcriptional programs from productive DNA synthesis and is accompanied by replication-associated DNA-damage signaling and checkpoint activation.

## DISCUSSION

Large-scale transcriptomic studies routinely identify thousands of cancer-associated lncRNAs, yet determining which candidates warrant functional investigation remains a major challenge. Here, we integrated complementary disease-associated evidence across colorectal, gastric, and esophageal cancers and prioritized *LINC01094* for functional study in CRC. Across 52 CRC cell lines, endogenous LINC01094 expression defined a transcriptional state that was strongly opposed by acute CRISPRi-mediated depletion in HCT116 cells. *LINC01094* depletion impaired DNA synthesis and was accompanied by replication-associated DNA-damage signaling and checkpoint activation. Together, these findings support a role for *LINC01094* in maintaining a CRC-associated transcriptional state and illustrate how multidimensional disease evidence can guide functional investigation of cancer-associated lncRNAs.

Existing cancer genomic resources and lncRNA prioritization strategies have integrated tumor expression, clinical, genomic, and network information in complementary ways (31–34). In our analysis, these evidence dimensions were kept separate because they capture different aspects of tumor biology and are subject to different technical limitations. Their outputs were therefore integrated only during candidate selection rather than combined into a composite score. Recurrent tumor upregulation and malignant epithelial detectability served as common disease-context requirements, whereas clinical and network analyses provided parallel forms of support. We also avoided sequential filtering because lack of support in a single dataset could reflect differences in statistical power, cohort composition, or transcript detection rather than lack of biological relevance. The individual analyses recovered several lncRNAs previously implicated in GI cancer, including *LINC00152*, *LUCAT1*, and *RHPN1-AS1* in the clinical analysis (35–37); *MIR31HG*, *SNHG16*, and *AGAP2-AS1* in the network analysis (38–40). Convergence across evidence dimensions further prioritized *MIAT* and *HOXA11-AS* (10, 41). Recovery of these established cancer-associated lncRNAs provided an internal check that the strategy enriched for disease-relevant signals rather than simply recapitulating tumor-upregulated transcripts. These measures nevertheless remain associative: clinical relationships are observational, recurrent expression can retain lineage effects, and co-expression does not establish causal regulation. Their value therefore lies in convergence of complementary evidence for candidate selection, extending previous cohort-based analyses of lncRNA dysregulation in GI cancer (42–44).

Bulk tumor profiles do not resolve the cellular source of lncRNA expression, and variation in malignant epithelial, stromal, immune, endothelial, and other cellular compartments can influence tumor-associated expression, clinical association, and co-expression networks (16, 17). Although deconvolution methods can estimate these contributions from bulk transcriptomes (45, 46), we used the TabulaTIME GI cancer single-cell-derived metacell resource to assess candidate expression directly within malignant epithelial populations (26). Malignant-versus-other differential expression and malignant epithelial detectability provided complementary information. Differential expression identifies preferential enrichment in malignant cells, whereas detectability establishes transcript presence despite the low abundance and sparse capture that complicate single-cell lncRNA analysis (6, 7, 27, 28). In our analysis, malignant-versus-other enrichment was strongly context-dependent, whereas malignant epithelial detectability extended across a broader range of GI cancer contexts. Neither measure alone establishes function: enrichment can remain cancer-type-specific, whereas detectability does not imply malignant-cell specificity or dependence. Considered together, these analyses add cellular context to bulk-derived associations without excluding candidates solely because they lack differential enrichment in a sparsely sampled single-cell dataset.

Previous studies have linked *LINC01094* to CRC progression, poor prognosis, proliferation, and invasion (20), and have proposed distinct mechanisms of action across tumor types (47–50). In HCT116 cells, we found that the predominant *LINC01094* transcript was enriched in the nucleus. By contrast, the previous CRC study that defined a miR-1266-5p/SLPI ceRNA model reported predominantly cytoplasmic *LINC01094* (20). Consistent with a distinct molecular context, *SLPI* showed only a modest, non-significant response in HCT116 RNA-seq and no detectable change by RT-qPCR following *LINC01094* depletion in either HCT116 or Caco-2 cells (Fig. S6C). Other studies have linked *LINC01094* to miR-340-5p/E2F3 signaling (47) and PKM2/JMJD5 complex recruitment in breast cancer (50), and to AZGP1-dependent PTEN/AKT signaling and RBMS2/HDAC1-dependent regulation of *CDKN1A* in gastric cancer (48, 49). In HCT116 cells, p21 accumulated following *LINC01094* depletion without a corresponding increase in *CDKN1A* mRNA, further distinguishing the response observed here from this gastric cancer model. The broader *LINC01094*-associated transcriptional architecture was likewise lineage dependent, with only partial recapitulation in STAD and a distinct profile in ESCA (Fig. S6E). Together, these observations indicate that *LINC01094* can engage distinct regulatory outputs across cellular and tumor contexts.

The induction of replication/checkpoint and DDR programs in HCT116 cells occurred despite reduced cell growth and DNA synthesis, indicating that this transcriptional response did not reflect productive cell-cycle activation. Increased γH2AX, CHK1 Ser345 phosphorylation, and p21, together with elevated γH2AX foci in residual EdU-positive cells, instead support replication-associated DNA-damage signaling and checkpoint activation. Apoptotic execution was not detected, and senescence-associated markers were discordant, arguing against a coordinated apoptotic or senescence response. These findings do not distinguish primary replication stress from impaired processing or repair of replication-associated DNA damage. Although the growth defect extended to Caco-2 cells, the weaker replication/checkpoint marker response in this model suggests that the magnitude or composition of the downstream response may vary across CRC cellular contexts.

Our data define a CRC-associated *LINC01094* transcriptional state across 52 cell lines and link *LINC01094* depletion to impaired growth across two CRC models and to broader transcriptional remodeling in HCT116 cells. In HCT116 cells, where *LINC01094-1* was the predominant nuclear-enriched transcript, the molecular basis of *LINC01094* activity remains unresolved. Recent work tracing *CRNDE* function to a defined isoform and discrete internal RNA element illustrates the importance of resolving lncRNA function below the locus level (51). Similar studies, including orthogonal perturbation approaches, will be needed to define the molecular activities of *LINC01094* and the additional candidates identified here and to determine how these functions vary across cellular contexts (52–54). Extending this framework across additional tumor types and increasingly cell-resolved or perturbational datasets will help distinguish lineage-restricted lncRNA signals from recurrent cancer-associated programs. More broadly, the same strategy can be adapted beyond cancer by replacing tumor-specific dimensions with disease-matched evidence and coupling these signals to functional perturbation to move from association toward functional prioritization.

## ACKNOWLEDGEMENTS

We thank Chelsea Denniss, Isabella DiLollo, and members of the Lee laboratory for experimental assistance and helpful suggestions on the manuscript. We thank Dr. Patricia Loomis and the Confocal Facility at Rosalind Franklin University of Medicine and Science (RFUMS) for assistance with fluorescence microscopy and imaging. We also acknowledge the patients and investigators who contributed to the publicly available datasets analyzed in this study. This study utilized data generated by the TCGA Research Network: https://www.cancer.gov/tcga. This work was supported by startup funds from RFUMS to J.-S.L.

## AUTHOR CONTRIBUTIONS

J.-S.L. conceived and supervised the study. B.F., P.L., E.P., J.W. and J.-S.L. designed experiments and interpreted the results. B.F. and E.P. performed experimental studies. B.F., P.L., and N.S. performed bioinformatic analyses. R.J.B. assisted with data interpretation and manuscript editing. J.W. provided technical assistance and critical resources. B.F. and J.-S.L. wrote the manuscript.

## COMPETING INTERESTS

The authors declare no competing interests.

## ETHICS STATEMENT

This study analyzed de-identified, publicly available datasets and established human cancer cell lines and did not involve new human subject recruitment or intervention.

## MATERIALS AND METHODS

### Bulk TCGA data acquisition and preprocessing

Gene-level raw counts and clinical metadata for TCGA colon adenocarcinoma (COAD), rectal adenocarcinoma (READ), stomach adenocarcinoma (STAD), and esophageal cancer (ESCA) were downloaded from the GDC Data Portal (55) (Release 45.0) with TCGAbiolinks (56). STAR-aligned counts against the GDC GRCh38 reference were used throughout. COAD and READ were combined as colorectal cancer (CRC). Gene biotypes were assigned from GENCODE Release 49 (GRCh38.p14) (57) by Ensembl gene identifier. Matched Primary Tumor and Solid Tissue Normal samples were identified by TCGA patient barcode, yielding 51 CRC, 32 STAD, and 13 ESCA pairs (Table S1A). Tumor-only analyses used 633 CRC, 443 STAD, and 185 ESCA Primary Tumor samples (Table S2A). All analyses were performed independently within each cancer type. To visualize *LINC01094* expression, TPM values from CRC paired tumor– normal samples (n = 51) and log2(TPM + 1) values from CRC tumor samples with evaluable T stage annotation (n = 627) were plotted.

### Principal component analysis and tumor–normal differential expression

PCA used DESeq2 variance-stabilized values for the top 500 most variable genes. Differential expression was performed separately for CRC, STAD, and ESCA with DESeq2 on raw counts (58), retaining genes with ≥10 counts in ≥2 samples per cohort. Paired comparisons used the design ∼ patient + condition with normal tissue as reference. Wald *P* values were adjusted by Benjamini–Hochberg, and log2FC were shrunken with apeglm (59) via lfcShrink. Genes were called differentially expressed at |shrunken log2FC| ≥ 0.5 and FDR < 0.05; tumor-upregulated lncRNAs were those meeting the same cutoffs in the positive direction. Cross-cancer concordance was assessed by pairwise Pearson correlation of shrunken log2FC, computed separately for protein-coding genes and lncRNAs using genes tested in both cohorts. Recurrent pan-GI activation was defined as upregulation in ≥2 of 3 cohorts. Overlaps were visualized with ComplexHeatmap UpSet plots (60) (Table S1).

### Clinical progression and survival analyses

Clinical analyses used Primary Tumor samples with available expression and clinicopathologic annotation. Raw counts were normalized by DESeq2 variance-stabilizing transformation (VST). AJCC pathological T, N, and M annotations were collapsed to major stage categories, treated as ordinal, and filtered to exclude Tx, Nx, and Mx. Stage-associated expression was tested within each cancer type and staging axis by DESeq2 likelihood ratio test on raw counts, comparing a full model containing the ordinal stage variable with an intercept-only reduced model. LncRNAs with nominal LRT *P* < 0.05 were considered stage-associated. Spearman rank correlation on VST values assigned directionality, and lncRNAs with LRT *P* < 0.05 and *r*s > 0 were retained for the directional progression layer. Overall survival was analyzed by univariate Cox proportional hazards regression on cohort-standardized VST values, with hazard ratios reported per one standard deviation increase. LncRNAs with hazard ratio > 1 and nominal Cox *P* < 0.05 were classified as poor-survival-associated. Kaplan–Meier curves used cohort-specific median stratification. Adjusted statistics are reported in Table S2.

### Weighted gene co-expression network analysis

WGCNA (19) was performed on TCGA Primary Tumor samples from CRC, STAD, and ESCA. Raw counts were processed with edgeR (61), normalized by trimmed mean of M-values, and converted to log2 counts per million. Protein-coding and noncoding genes with CPM > 1 in ≥20% of samples were retained. Genes and samples with excessive missingness or zero variance were removed with WGCNA quality-control functions. A pooled network was built from the integrated three-cohort matrix. Soft-thresholding power was evaluated with pickSoftThreshold and set to 10 based on scale-free topology fit while retaining connectivity.

Modules were detected with blockwiseModules using power = 10, networkType = “unsigned”, TOMType = “unsigned”, corType = “pearson”, minModuleSize = 30, mergeCutHeight = 0.25, deepSplit = 2, maxBlockSize = 5000, pamStage = TRUE, and pamRespectsDendro = FALSE. The grey unassigned set was excluded from module prioritization. Module eigengenes were correlated by Pearson with binary CRC, STAD, and ESCA indicators and with harmonized ordinal AJCC T, N, and M variables. The maximum absolute module–trait correlation summarized trait association for each module is in Table S3.

### Module prioritization, lncRNA ranking, and preservation analysis

Pooled modules were evaluated on three features: module–trait association strength (maximum absolute eigengene–trait Pearson correlation), lncRNA representation (fraction of module genes annotated as lncRNAs), and lncRNA module membership, quantified as eigengene-based connectivity (kME), the correlation between each lncRNA and the eigengene of its assigned module. Network-prioritized modules were non-grey modules with at least one significant module–trait association (Pearson FDR < 0.05) and at least one lncRNA with kME ≥ 0.5, yielding 16 of 21 non-grey modules. LncRNAs were considered network supported if they were assigned to a network-prioritized module. GO-BP enrichment for prioritized modules was performed with clusterProfiler (62), mapping Ensembl to Entrez identifiers with org.Hs.eg.db. All WGCNA-tested genes were used as background and Benjamini–Hochberg FDR < 0.05. Module preservation across CRC, STAD, and ESCA was assessed with modulePreservation (200 permutations), interpreting Zsummary < 2, 2–10, and > 10 as no, weak-to-moderate, and strong preservation, respectively. Modules with Zsummary > 10 in all three cohorts were classified as strongly preserved across all three GI cancer cohorts. Modules with strong preservation in at least one cohort and weak-to-moderate preservation in at least one other were classified as cohort-influenced. Cancer-type-specific networks were built independently for each cohort using the same preprocessing and module-detection framework, with soft-thresholding powers selected separately. LncRNAs in prioritized pooled modules were ranked by absolute pooled kME. Cancer-type-resolved support was annotated by comparing module assignment and kME across the three cohort-specific networks (Table S3).

### Single-cell-derived metacell analysis

Metacell profiles and metadata were obtained from the TabulaTIME pan-cancer single-cell atlas (26) (accessed December 10, 2025). Colorectal, gastric, and esophageal cohorts were extracted using the atlas cancer-type field and analyzed as CRC, STAD, and ESCA. Major cell-type annotations were taken from the curated atlas field. Profiles annotated as malignant defined the malignant epithelial compartment, and all other annotated profiles were assigned to non-malignant compartments. Precomputed atlas UMAP coordinates were used for visualization. Annotations were supported with canonical markers for epithelium (*EPCAM*, *KRT8*, *KRT18*), immune cells (*PTPRC*, *CD3D*, *CD3E*, *MS4A1*, *LYZ*), fibroblasts and stroma (*COL1A1*, *COL1A2*, *ACTA2*), endothelium (*PECAM1*, *VWF*), and proliferation (*MKI67*, *TOP2A*). Profile counts, patient sample counts, and compartment annotations are provided in Table S4A.

### Pseudobulk malignant-versus-other differential expression

TabulaTIME normalized values were aggregated by patient sample and cell group for pseudobulk analysis. Only samples containing both malignant and non-malignant profiles were retained, yielding 32 CRC, 52 ESCA, and 25 STAD samples. Aggregated values were transformed as log2(mean normalized expression + 1), and differential expression was performed within each cohort using limma (63) with donor effects included in the design matrix and empirical Bayes moderation (eBayes). Malignant-associated lncRNAs were defined by positive malignant-versus-other log2FC and Benjamini–Hochberg adjusted *P* < 0.05, with no minimum fold-change threshold (Table S4B).

### Detection-based scoring and single-cell candidate prioritization

For detection-based scoring, the fraction of profiles with detectable expression and the average normalized expression were computed for each lncRNA within each cancer context. Transcripts detected in ≥1% of malignant profiles were classified as detection-supported, and detection-supported status was compared across CRC, STAD, and ESCA. The complete scoring matrix is provided in Table S4B. For the cell-type dot plot, the top ten unique malignant-associated candidates were selected by maximum malignant-versus-other log2FC across the three contexts. For cohort-level profiling, the top 40 unique detection-supported lncRNAs from the malignant-associated subset were selected by the same ranking. Cohort-average expression was Z-scored by row and clustered with pheatmap (Euclidean distance, complete linkage), with cohorts held in fixed column order.

### Human cell culture

HCT116 cells (male), Caco-2 cells (male), and HEK293T cells (female) were cultured in Dulbecco’s Modified Eagle Medium (DMEM) (Fisher Scientific, MT10013CV) supplemented with 10% fetal bovine serum (Fisher Scientific, A5256701) and 1% penicillin/streptomycin (Fisher Scientific, MT30002CI). Cell lines were confirmed to be free of mycoplasma contamination.

### Generation of CRISPRi cell line

HCT116 cells and Caco-2 cells were transduced with a dCas9^KRAB^-mCherry lentivirus (Addgene #60954) (64), and a pooled population stably expressing high levels of mCherry was established by fluorescence-activated cell sorting (FACS) of the top 30% of mCherry-positive cells. HEK293T cells were used for lentiviral production.

### CRISPRi-mediated gene knockdown

sgRNAs were designed using a previously described algorithm (65). Two sgRNAs targeting the *LINC01094* TSS and one non-targeting control were designed. Sequences are provided in Table S5C. Individual sgRNAs were cloned into pU6-sgRNA lentiviral vector (Addgene #60955) as previously described (64). HCT116 CRISPRi cells were transduced with lentiviral vectors sgNT, *LINC01094-*sg1, and *LINC01094-*sg2 in medium containing 8 μg/mL polybrene (Santa Cruz Biotechnology, SC-134220). Caco-2 CRISPRi cells were transduced with lentiviral vectors sgNT and *LINC01094-*sg1 in medium containing 8 μg/mL polybrene. After transduction, cells were cultured for 1 day, followed by selection in 2 μg/mL puromycin for 3 days prior to analysis. For selected follow-up assays, *LINC01094*-sg1 was used unless otherwise specified because it produced greater knockdown than sg2.

### Cumulative cell growth assay

HCT116 cells were seeded at 2.5 x 10^5^ cells per well in 6-well plates containing 2 mL of medium. Cells were counted and passaged every 2 days, with 2.5 x 10^5^ cells re-seeded per well after each passage. Caco-2 cells were seeded at 1.0 x 10^5^ cells per well in 6-well plates containing 2 mL of medium. Cells were counted and passaged every 3 days, with 1.0 x 10^5^ cells re-seeded per well after each passage. Cell numbers were determined at each passage using a TC20 automated cell counter (Bio-Rad, 1450102), and cumulative cell numbers were calculated across successive passages. All growth assays were performed with three biological replicates.

### RNA isolation and quantitative RT-PCR (RT-qPCR)

Total RNA was extracted from cells using TRIzol followed by RNA Clean & Concentrator-5 (Zymo Research, R1014) with on-column DNase digestion. cDNA was synthesized from 1 μg of total RNA using the PrimeScript RT Master Mix (Takara, RR036A) according to the manufacturer’s instructions. SYBR Green PCR Master Mix (Bio-Rad, 1725271) was used for qPCR reactions. RNA expression levels were normalized to *GAPDH* using the ΔΔCt method. Primer sequences used for RT-qPCR are provided in Table S5C.

### Endpoint RT-PCR

cDNA was synthesized from 1 μg total RNA using the PrimeScript RT Master Mix (Takara) according to the manufacturer’s instructions and subsequently amplified using Q5 HotStart Master Mix (NEB). Primer sequences used for endpoint RT-PCR are provided in Table S5C.

### Subcellular fractionation

Cytoplasmic and nuclear fractions were prepared as described previously (66, 67). Cells were harvested by trypsinization and lysed in 250 μL cytoplasmic lysis buffer (50 mM Tris-HCl pH 8.0, 140 mM NaCl, 1.5 mM MgCl2, 0.5% NP-40, 0.2 U/μL SUPERaseIn (Thermo Fisher, AM2696)), incubated 5 min on ice, and centrifuged at 1,300 x *g* for 5 min at 4 °C. The cytoplasmic supernatant was cleared at 16,000 x *g* for 15 min at 4 °C. The nuclear pellet was washed in cytoplasmic lysis buffer to remove residual cytoplasm and resuspended in 250 μL nuclear lysis buffer (50 mM Tris-HCl pH 8.0, 500 mM NaCl, 1.5 mM MgCl2, 0.5% NP-40, 0.2 U/μL SUPERaseIn). RNA was extracted from each fraction with TRIzol followed by RNA Clean & Concentrator-5 (Zymo Research, R1014) with on-column DNase digestion. RT-qPCR primers are listed in Table S5C.

### Antibodies

The following antibodies were used in this study: PCNA (rabbit polyclonal, Proteintech, 10205-2-AP), p21 (rabbit polyclonal, Proteintech, 10355-1-AP), γH2AX (rabbit monoclonal, Proteintech, 83307-2-RR), β-actin (rabbit polyclonal, Proteintech, 20536-1-AP), p-CHK1 (Ser345) (rabbit polyclonal, Proteintech, 28803-1-AP), total CHK1 (mouse monoclonal, Proteintech, 60277-1-Ig), cleaved caspase-3 (rabbit polyclonal, Proteintech, 25128-1-AP), GAPDH (rabbit monoclonal, Cell Signaling Technology, 14C10), Multi-rAb CoraLite Plus 750-Goat Anti-Rabbit Recombinant Secondary Antibody (Proteintech, RGAR006), Multi-rAb CoraLite Plus 488-Goat Anti-Rabbit Recombinant Secondary Antibody (Proteintech, RGAR002), Multi-rAb CoraLite Plus 647-Goat Anti-Rabbit Recombinant Secondary Antibody (Proteintech, RGAR005), Multi-rAb CoraLite Plus 750-Goat Anti-Mouse Recombinant Secondary Antibody (Proteintech, RGAM006). Antibodies were used at a dilution of 1:1000 in 2% BSA for western blotting and 1:250 in 5% BSA for immunofluorescence.

### Western blotting

Cells were lysed in 2× NuPAGE LDS Sample Buffer (Thermo Fisher Scientific, NP0007) for 5 min at room temperature followed by 5 min at 95 °C. sgNT HCT116 cells used as the positive control for caspase-3 cleavage were treated with 1 μM doxorubicin for 24 hours before lysis. Proteins were separated on 4–12% Bis-Tris NuPAGE gels and transferred to nitrocellulose membranes (0.45 μm, Fisher Scientific, 45004002). Membranes were blocked in TBST with 5% BSA for 30 min at room temperature, incubated with primary antibodies overnight at 4 °C, then with IR dye-labeled secondary antibodies for 1 h at room temperature. Blots were imaged on a LI-COR Odyssey and band intensities quantified in ImageJ/Fiji. Membranes were physically sectioned by molecular-weight region before antibody incubation. Uncropped original membrane sections are provided in the Supplementary Information.

### Long-read sequencing analysis

Long-read RNA-seq data obtained from the NCBI Sequencing Read Archive under accession numbers PRJNA727942 (HGC-27) and PRJNA1282689 (AGS) were aligned to GRCh38/hg38 with minimap2 (68) (v2.26) using the splice-aware setting (-ax splice -uf -k14) against an index built with minimap2 -d. Alignments were converted to BAM, sorted, and indexed with samtools (69) (v1.19.2) for genomic visualization.

### RNA sequencing (RNA-seq)

RNA was extracted with TRIzol followed by RNA Clean & Concentrator-5 (Zymo Research, R1014). Total RNA was submitted to the Plasmidsaurus 3′-end counting RNA-seq service, which performs reverse transcription and second-strand synthesis followed by tagmentation, indexing, and PCR amplification to generate 3′-biased libraries. Sequencing was performed on an Illumina NovaSeq X Plus.

### RNA-seq data analysis

Read quality was assessed with FastQC. Reads were filtered with fastp using poly-X tail trimming, 3′ quality trimming, minimum Phred 15, and minimum length 50 bp (70), then aligned with STAR with non-canonical junction removal and unmapped-read output (71), and coordinate-sorted with samtools (69). PCR and optical duplicates were removed by UMI-based deduplication with UMIcollapse (72). Alignment metrics, strand specificity, and feature distribution were assessed with RSeQC and Qualimap and aggregated with MultiQC (73–75). Expression was quantified with featureCounts using strand-specific counting, fractional assignment of multi-mapping reads, exons and 3′ UTRs as features, and grouping by gene ID; counts were annotated with gene biotype from the reference GTF. Sample–sample Pearson correlations for heatmaps and PCA used TMM-normalized counts. Differential expression was assessed with edgePython, a Python implementation of edgeR (61). For the acute-depletion analysis, cells expressing *LINC01094-*sg1 and *LINC01094-*sg2 were combined as the depletion group and compared with sgNT cells. Genes were ranked by moderated t-statistic, and GSEA was performed with clusterProfiler against GO-BP gene sets (62). Pathways with Benjamini-Hochberg FDR < 0.05 were considered significantly enriched.

### CCLE acquisition and continuous expression modeling

CCLE transcriptomic profiles were retrieved from EBI (E-MTAB-2770) as TPM values. Missing values were set to 0 TPM and expression was transformed as log2(TPM + 1).

Cell lines annotated “colon,” “rectal,” or “colorectal” defined the CRC cohort, and genes with TPM > 0.1 in at least three CRC lines were retained. Gene-wise linear models were fitted with limma using continuous log2(TPM + 1) *LINC01094* expression as the predictor, with empirical Bayes moderation. Genes were ranked by moderated *t*-statistic and GSEA was performed with clusterProfiler against GO-BP gene sets, mapping Ensembl to Entrez identifiers with org.Hs.eg.db. Pathways with Benjamini–Hochberg FDR < 0.05 were considered significantly enriched. To compare the acute-depletion enrichment with the CCLE analysis, GO terms FDR-significant in both analyses were considered shared, and terms that were FDR-significant in both analyses and had opposite-signed NES in each analysis were considered reciprocal. NES correlations were calculated using both Spearman and Pearson correlation tests.

### Sensitivity analysis of CRC-restricted pathway signatures

CRC cell lines were reclassified as *LINC01094*-high or -low using either a CRC-specific median threshold or a fixed cutoff (TPM > 0.2 versus ≤ 0.2). Differential expression was performed with limma and empirical Bayes moderation, genes were ranked by moderated *t*-statistic, and GSEA was performed as above. Enrichment direction and NES were compared with the primary analysis to test whether the major programs were reproduced.

### Lineage-dependent gene set enrichment analyses

Cell lines annotated “esophagus” or “esophageal” comprised the ESCA cohort and those annotated “stomach” or “gastric” the STAD cohort. Preprocessing, expression filtering, and transformation were as for CRC. Within each lineage, cell lines were stratified by lineage-specific median *LINC01094* TPM, and differential expression and GSEA were performed as above.

### EdU incorporation assay

HCT116 cells were cultured for 24 h. Medium was replaced with fresh medium containing 10 μM EdU, and cells were incubated for 30 min. sgNT positive-control cells shown in Fig. S6F were treated with 1 μM doxorubicin for 16 h before EdU labeling. Cells were fixed in 3.7% formaldehyde, permeabilized with 0.5% Triton X-100, and labeled by click chemistry for 30 min in 50 mM Tris pH 7.5, 150 mM NaCl, 5 mM sodium ascorbate, 1 mM copper (II) sulfate, 5 μM azide-fluor 488 (MilliporeSigma, 760765-1MG). Cells were counterstained with DAPI and imaged on a ZOE Fluorescent Cell Imager (Bio-Rad, 1450031). Experiments were performed three times with approximately 200 cells counted per replicate, and images were analyzed in ImageJ.

### Immunofluorescence

EdU labeling and click-chemistry detection were performed as described above. Cells were then blocked in 5% BSA/PBS for 30 min at room temperature, incubated with primary antibody overnight at 4 °C, and with secondary antibody for 1 h at room temperature. Coverslips were mounted with SlowFade Diamond Antifade Mountant (Fisher Scientific, S36967). Images were acquired on an Olympus FV10i confocal microscope with a 63× oil objective. Experiments were performed three times with approximately 100 cells counted per replicate, and images were analyzed in ImageJ.

### Visualization and statistical analysis

Plots were generated in R with ggplot2, ComplexHeatmap (60), pheatmap (76), survminer (77), and related packages; plot types and the quantities displayed are specified in the figure legends. Unless otherwise stated, multiple-testing correction used the Benjamini–Hochberg method. For exploratory prioritization layers, including clinical progression and survival analyses, nominal *P* value thresholds were used for candidate retention, with adjusted *P* values reported in the corresponding supplementary tables to permit evaluation at alternative stringency. All experiments were performed with at least three independent biological replicates as indicated in each figure legend, values are reported as mean ± SD, and statistical significance was assessed in Prism 11 (GraphPad Software).

### Software and reproducibility

Analyses were performed in R version 4.5.1 using Bioconductor and CRAN packages including TCGAbiolinks v2.36.0, SummarizedExperiment v1.28.1, DESeq2 v1.48.2, apeglm v1.30.0, edgeR v4.6.3, limma v3.64.3, WGCNA v1.73, clusterProfiler v4.16.0, org.Hs.eg.db v3.21.0, survival v3.8.3, survminer v0.5.1, ComplexHeatmap v2.24.1, pheatmap v1.0.13, and ggplot2 v4.0.1. Command-line analyses used minimap2 v2.26, FastQC v0.12.1, fastp v0.24.0, STAR aligner v2.7.11, samtools v1.22.1, UMIcollapse v1.1.0, RSeQC v5.0.4, Qualimap v2.3, MultiQC v1.32, featureCounts v2.1.1, edgePython v0.2.5. Random seed was set to 12345 for stochastic or permutation-based analyses where applicable, including WGCNA module preservation testing.

### Data availability

The RNA-seq data generated for this study have been deposited in the Gene Expression Omnibus (GEO) under accession number GSE334476. Reviewers may access this data using the token sbqnwycobnehpif. Long-read sequencing data analyzed in this study were obtained from the NCBI Sequencing Read Archive (SRA) under accession numbers PRJNA727942 and PRJNA1282689.

## SUPPLEMENTARY FIGURE LEGENDS

**Figure S1. Pan-GI cancers exhibit recurrent tumor-associated transcriptional remodeling and lncRNA dysregulation**

(A) Cancer-type-specific PCA of variance-stabilized expression profiles from CRC, STAD, and ESCA tumor and matched normal samples.

(B) Tumor–normal differential expression patterns of select housekeeping, glycolytic/metabolic, cell-cycle/proliferation, and ribosome/translation-associated marker genes. Dots show shrunken log2FC and –log10 FDR from DESeq2 analyses.

**Figure S2. Clinical progression and survival analyses prioritize recurrent lncRNA candidates across GI cancers**

(A) Heatmap of Spearman rank correlation coefficients for lncRNAs evaluated across AJCC T, N, and M stages in CRC, ESCA, and STAD. Positive values indicate increased expression with advancing stage.

(B) Heatmaps showing representative N stage-ordered expression patterns of stage-associated lncRNAs in CRC, ESCA, and STAD. Samples are ordered by AJCC N stage, and expression values are shown after normalization and row scaling. Genes shown were filtered by LRT *P* < 0.05 and ranked by positive Spearman correlation with stage. Because CRC lacked N3 samples, only N0–N2 are shown.

(C) UpSet plots showing overlap between LRT-supported stage-associated lncRNAs and lncRNAs with positive Spearman directionality (Spearman *rs* > 0 and LRT *P* < 0.05), identifying candidates with both stage-associated variation and progression-linked expression patterns. M1 sample sizes were CRC, n = 89; STAD, n = 30; ESCA, n = 9 (Table S2B).

**Figure S3. Pan-GI co-expression and cancer-type-resolved network analyses define disease-associated transcriptional modules**

(A) Soft-thresholding power analysis for the pooled pan-GI WGCNA network, showing scale-free topology fit and mean connectivity across candidate powers. The selected power is indicated.

(B) Sample clustering analysis performed before pooled network construction to assess sample-level relationships.

(C) Module size distribution for the pooled pan-GI WGCNA network.

(D–F) Cancer-type-resolved module–trait correlation heatmaps for CRC (D), STAD (E), and ESCA (F). Module eigengenes were correlated with available clinical variables within each cohort.

**Figure S4. Single-cell RNA-seq analyses reveal context-dependent malignant epithelial lncRNA expression across GI cancers**

(A) Integrated UMAP of the GI cancer subset colored by cancer type, showing the contribution of CRC, STAD, and ESCA profiles to the integrated single-cell-derived landscape.

(B) Dot plot of canonical marker genes used to support major cell-type annotation, including epithelial, immune, fibroblast/stromal, endothelial, and proliferation markers. Dot size indicates the fraction of profiles expressing each marker, and color indicates average normalized expression.

**Figure S5. *LINC01094* expression and transcript characterization**

(A) TCGA RNA-seq analysis of *LINC01094* expression in paired CRC tumor and adjacent normal tissue samples. Expression is shown as TPM. n = 51 paired samples. *P* = 0.0265, two-sided paired t-test.

(B) *LINC01094* expression across AJCC primary tumor T-stage categories in CRC tumors. Expression is shown as log2(TPM + 1). n = 627 tumors with evaluable T-stage annotation. Spearman *rs* = 0.16; *P* = 9.88E-05.

(C) IGV snapshot of publicly available long-read sequencing data from two gastric cancer cell lines at the *LINC01094* locus shows two alternatively spliced isoforms of *LINC01094*. Isoform information is summarized in Table S5B.

(D) *LINC01094* transcript models and RT-PCR validation in HCT116 cells. Top, schematic of the predicted *LINC01094* isoforms depicting the locations of CRISPRi-targeting sequence sg1 and RT-PCR primers (F1, R1, R2). Bottom, RT-PCR validation using two independent primer configurations. The F1 + R1 primer set (left), designed to amplify both predicted isoforms, detected a predominant, CRISPRi-sensitive amplicon consistent with the expected size range. The F1 + R2 primer set (right) specifically amplified *LINC01094-*1 at the expected size of 177 bp. -RT indicates the no-reverse-transcriptase control.

(E) Isoform-directed RT-qPCR quantification of the predicted *LINC01094* transcripts in HCT116 cells. Expression levels were normalized to *GAPDH*, and relative abundance is plotted with *LINC01094-1* set to 1. Data are presented as mean ± SD (n=3 biological replicates). \**P* <0.05; \*\**P* <0.01, calculated by two-tailed t-test.

(F) Distribution of endogenous *LINC01094* expression across the 52 CRC cell lines used for CRC-restricted CCLE modeling. Missing or non-detected TPM values were set to zero; HCT116 is indicated.

**Figure S6. Acute *LINC01094* depletion opposes transcriptional programs associated with endogenous *LINC01094* expression across CRC cell lines**

(A) PCA of HCT116 RNA-seq samples following CRISPRi-mediated *LINC01094* depletion with two independent sgRNAs compared with sgNT.

(B) RT-qPCR validation of representative RNA-seq marker genes following *LINC01094* depletion in Caco-2. Bar graph shows mean relative expression normalized to sgNT cells from three biological replicates. Seven selected HCT116-responsive markers showed directionally concordant changes in Caco-2 cells, with *TGM2* and *FN1* significantly reduced. \**P* <0.05, calculated by two-tailed t-test.

(C) RT-qPCR quantification of *SLPI* expression in HCT116 cells expressing sgNT, *LINC01094*-sg1, or *LINC01094*-sg2 and Caco-2 cells expressing sgNT or *LINC01094*-sg1. Expression levels were normalized to *GAPDH* and plotted relative to sgNT within each cell line. Data are presented as mean ± SD (3 biological replicates).

(D) Sensitivity analyses for the CRC-restricted CCLE model using threshold-stratified and median-stratified *LINC01094* expression groups. Representative GO-BP terms show that the major directional pathway patterns were preserved across stratification approaches.

(E) Cancer-type-restricted CCLE analyses in STAD and ESCA cell lines. Representative GO-BP terms show lineage-dependent *LINC01094*-associated pathway patterns across GI cancer contexts.

(F) Immunofluorescence of γH2AX in doxorubicin-treated HCT116 cells. Scale bar shows 10 μm.

## Notes

### Competing Interest Statement

The authors have declared no competing interest.

### Summary of Updates

This revised version substantially expands the original pan-gastrointestinal transcriptomic analysis with functional characterization of LINC01094 in colorectal cancer. We have incorporated analyses linking endogenous LINC01094 expression across 52 colorectal cancer cell lines to transcriptional responses following CRISPRi-mediated depletion. Functional validation has been expanded to include two colorectal cancer cell models, HCT116 and Caco-2. Additional experiments characterize impaired cell growth, reduced DNA synthesis, DNA-damage signaling, and checkpoint activation, including gamma-H2AX, CHK1 Ser345 phosphorylation, and p21 accumulation. We also examined LINC01094 subcellular localization and the previously reported SLPI-associated pathway. The manuscript has been substantially revised to incorporate these findings, with updated figures, methods, and supplementary materials.

