## Supplementary figures and images for "Pan-gastrointestinal prioritization links *LINC01094* to a colorectal cancer transcriptional state and replication-associated checkpoint activation"

### Supplemental figures

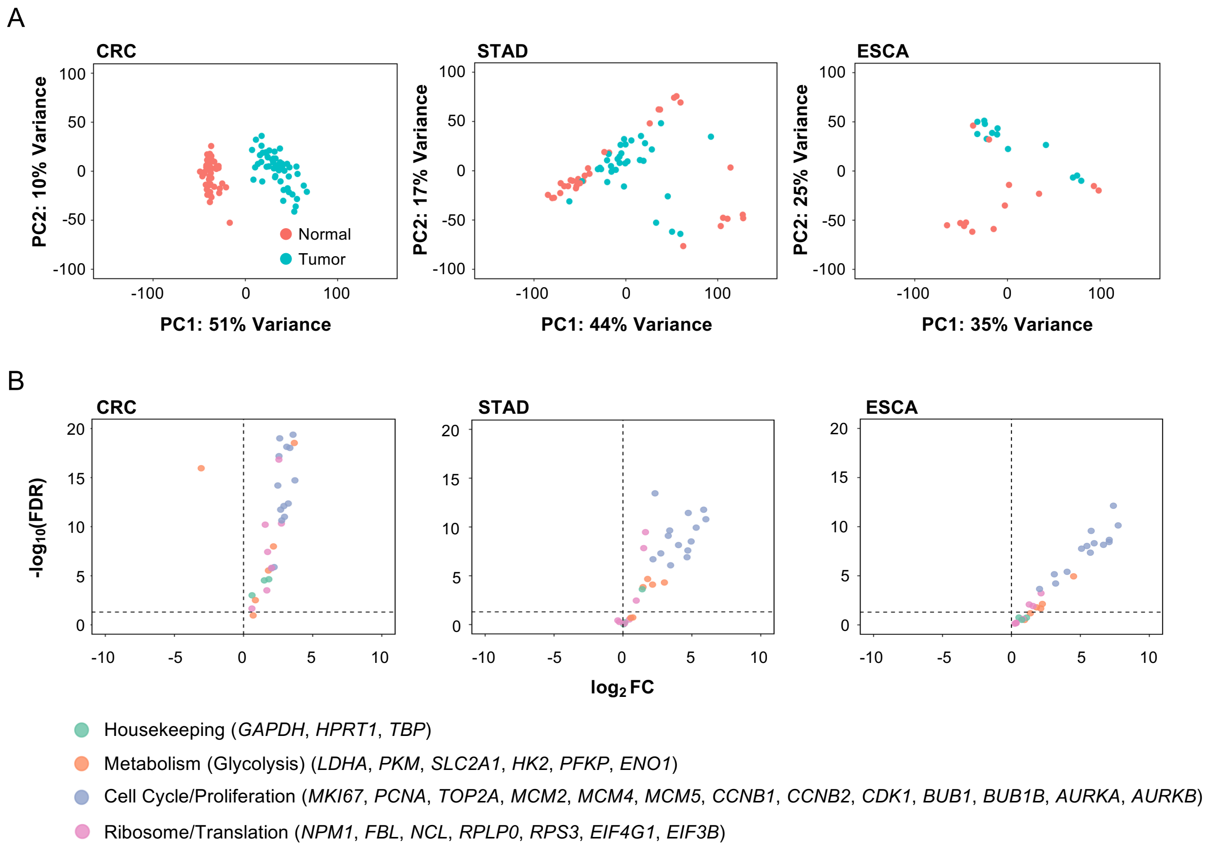

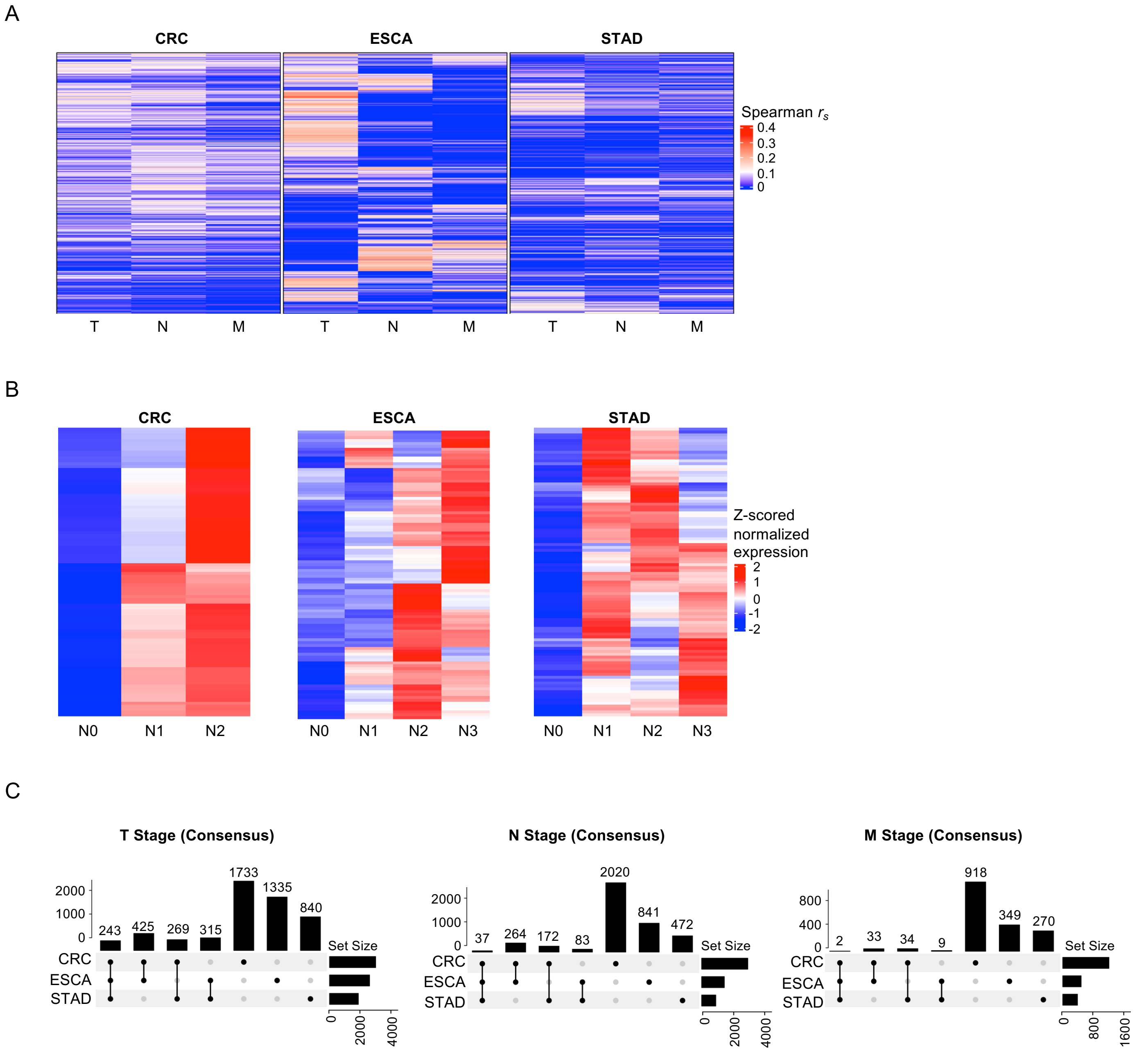

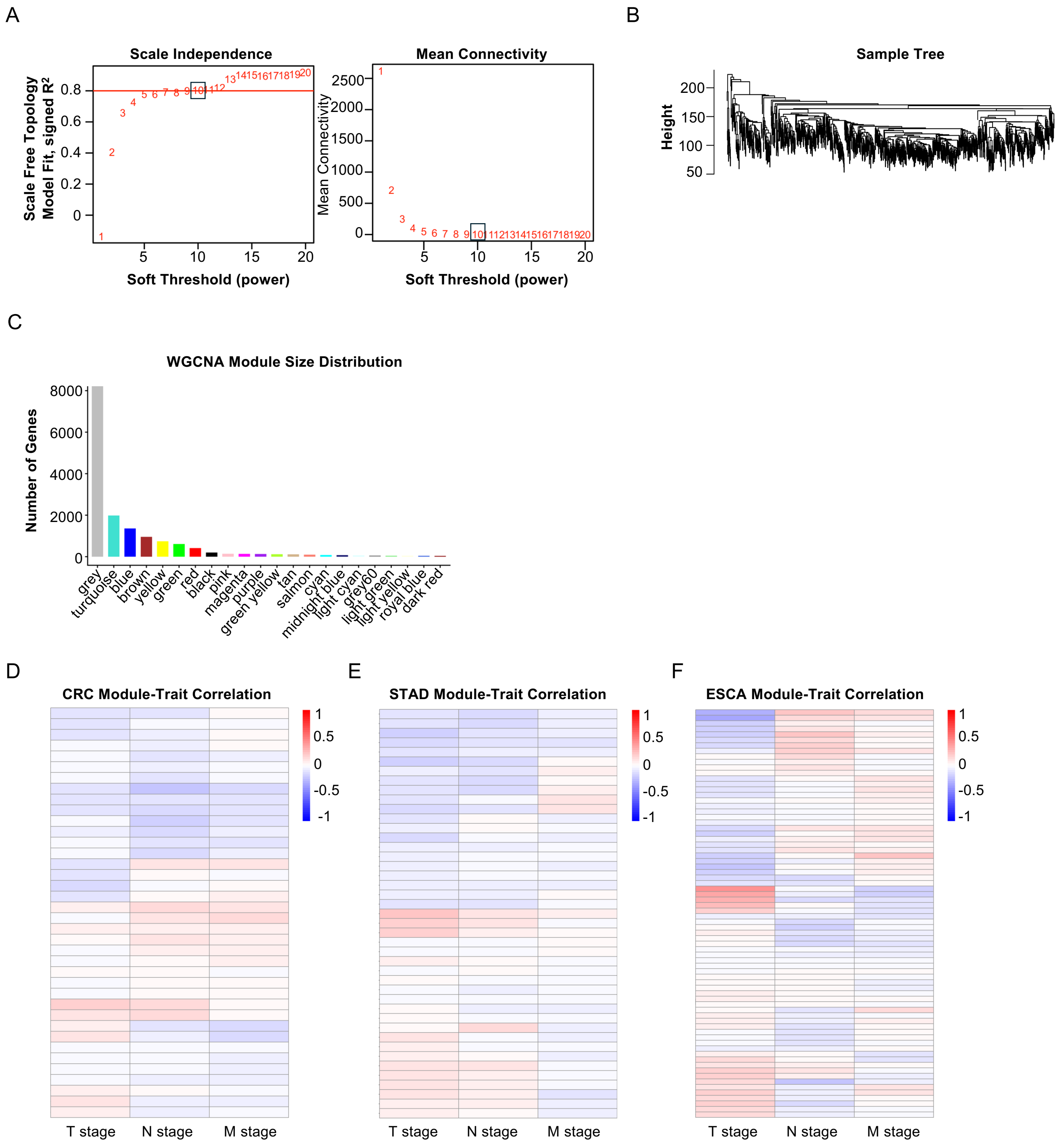

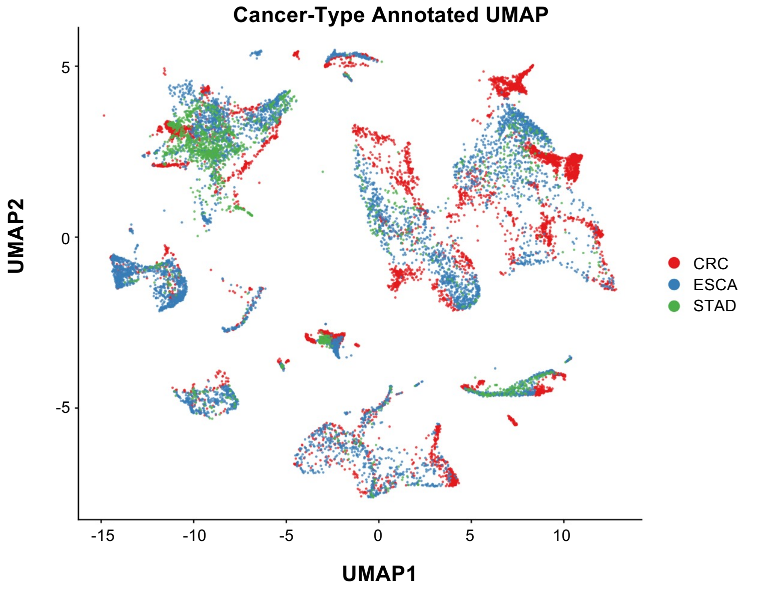

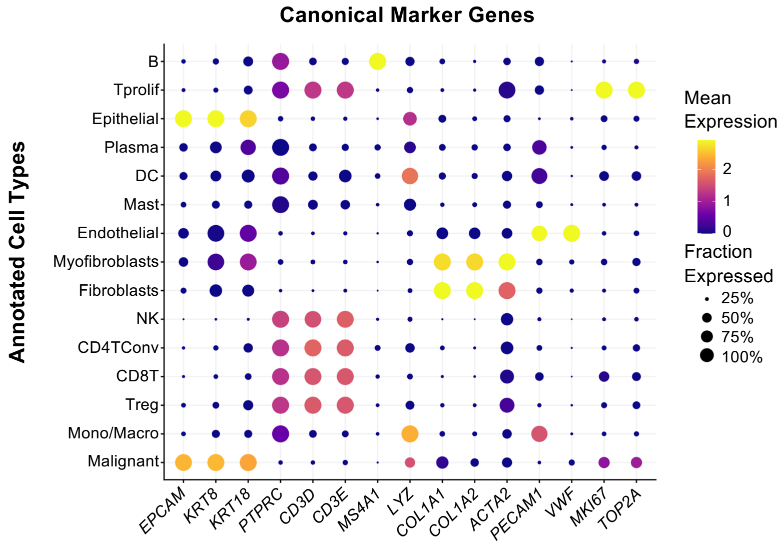

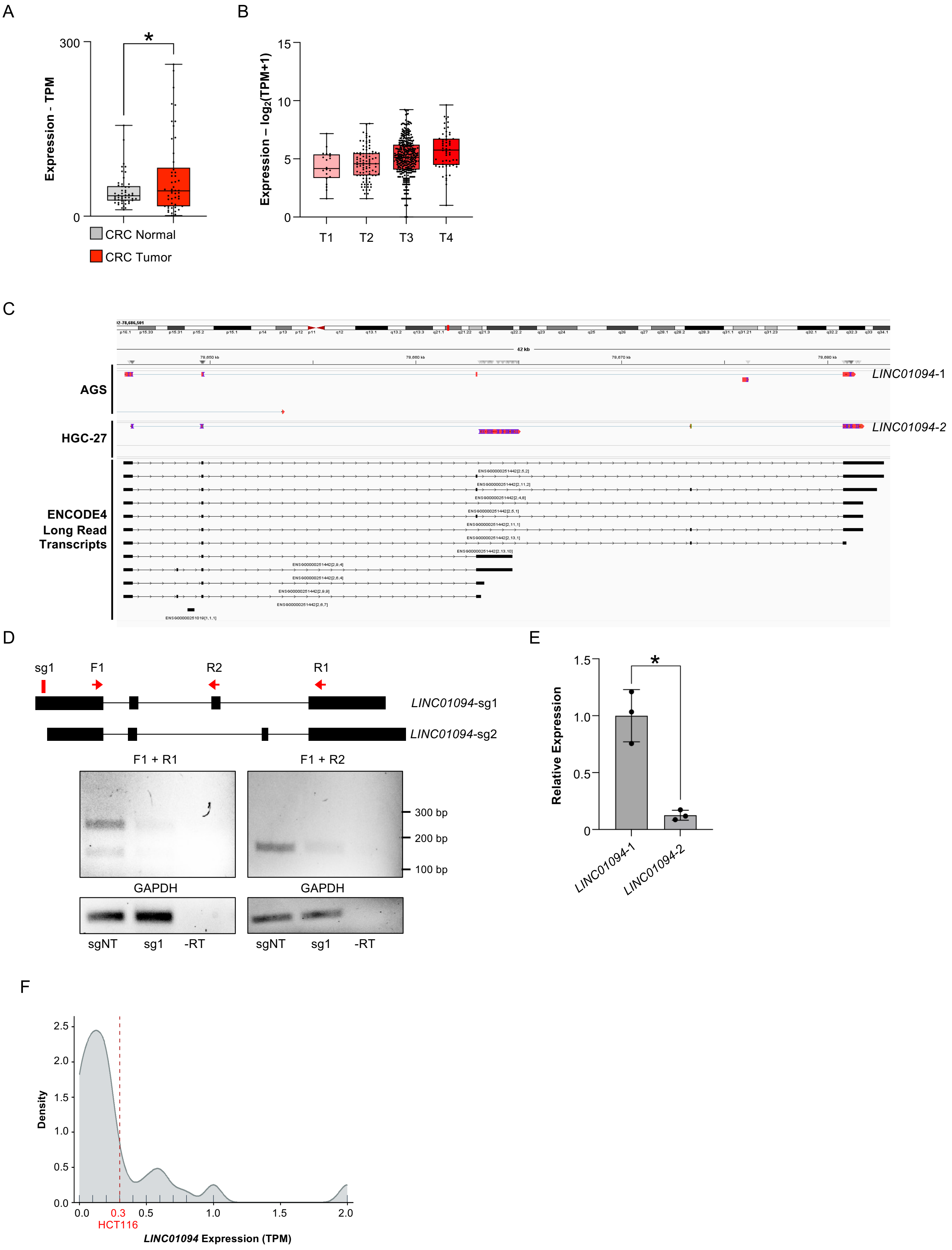

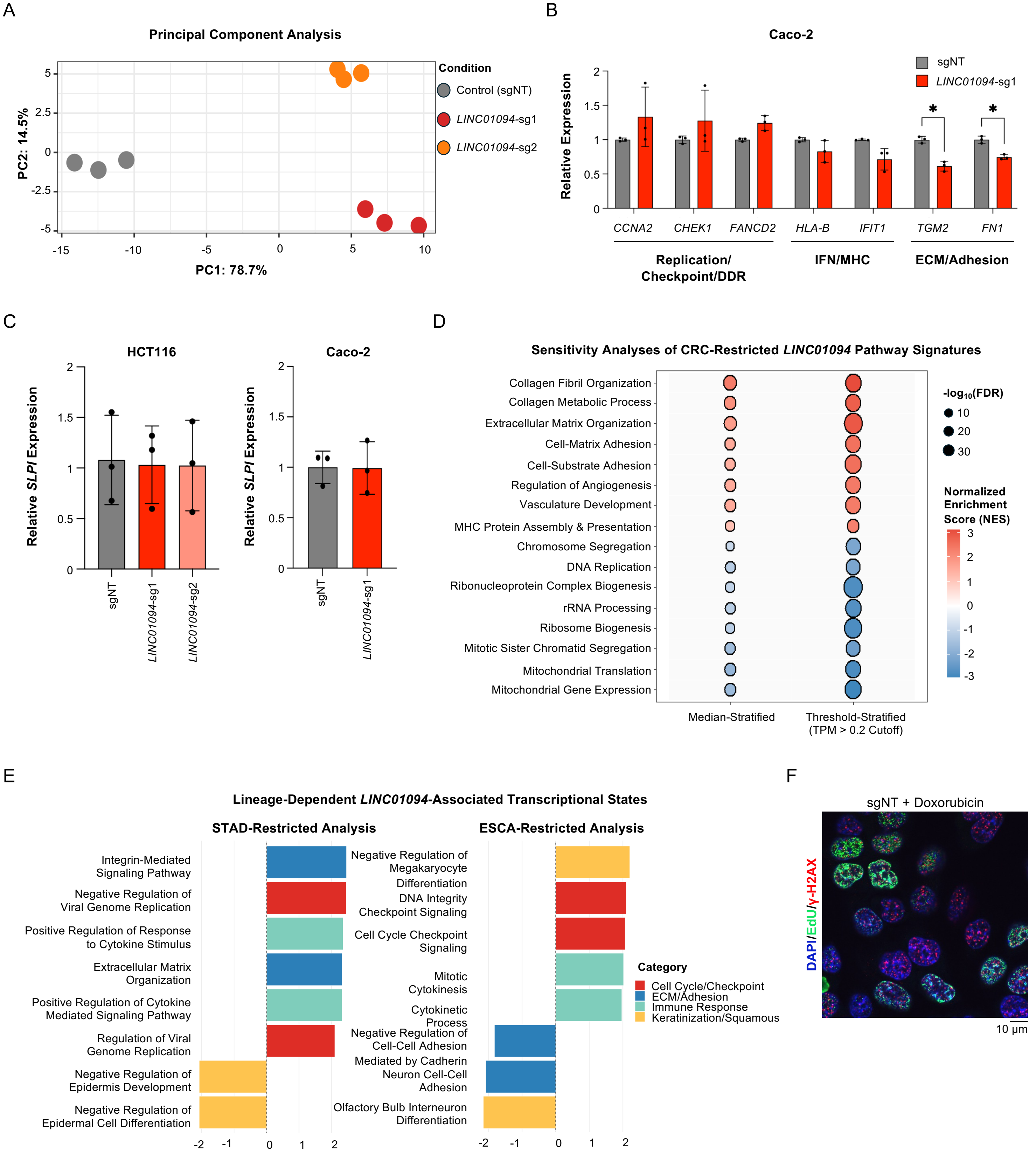
